# Modelling Maternal-to-Zygotic Transition Dynamics to Identify Common Gene Expression Baselines in Zebrafish Development

**DOI:** 10.64898/2026.11.17.767246

**Authors:** Marie gr. Holthaus, Paul Michaelis, Andreas Focks, Florian Schunck

## Abstract

We propose a semi-mechanistic ODE-based transcription model for zebrafish embryogenesis that captures common baseline expression dynamics by integrating the maternal-to-zygotic transition mechanism. We fitted this model to a time-course RNA-seq dataset of *D. rerio* spanning early developmental stages until 120 hpf. Using the inferred model parameters, we identified common gene expression pattern using fuzzy *c*-means clustering and compared these with clusters stemming from a shape-based approach using smooth trajectories. We show that the ODE-parameter-based approach improves to biological interpretability by linking clusters to underlying regulatory mechanism, such as the timing of zygotic genome activation, thereby generating more precise expression clusters. However, the results also revealed technical biases of the time-course data, originating from the poly(A) enrichment step.

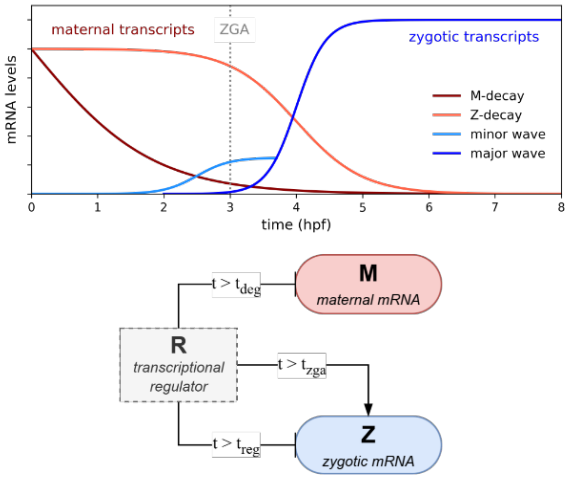

## 1 Introduction

Among vertebrates, the zebrafish (*Danio rerio*) is widely used as a model organism in biomedical and toxicology research. Several characteristics contribute to their popularity in the lab. The development progresses rapidly: the basic body plan is established within 24 hours post-fertilisation (hpf), and hatching occurs after 2 to 3 days post-fertilisation (dpf) [1, 2]. The zebrafish genome is fully sequenced [3], making zebrafish a powerful model for studying developmental gene expression and regulation. The embryo’s body is nearly transparent, which, for example, facilitates the observation of developmental abnormalities when exposed to chemicals or environmental stressors [4]. These attributes have promoted its use for estimating xenobiotic toxicity in fish [5, 6, 7]. More recently molecular toxicology has moved into the focus of *D. rerio* toxicity testing [8, 9, 10] to identify molecular triggers for chemical effects.

One approach is to compare gene expression levels between exposed and unexposed organisms, which, together with observations of lethal and sublethal effects, indicates the risk posed by the chemical at relevant environmental concentrations [11, 12, 13, 14].

However, during the early stages of embryonic development, gene expression is highly dynamic. In all animal embryos, the initial stages of development are driven exclusively by maternal gene products that are deposited into the egg during oogenesis [15, 16]. Those maternal mRNAs and proteins implement basic processes, direct the first cell divisions, and specify initial cell fate and patterning [17]. Meanwhile, the embryo is transcriptionally silent, unable to produce its own gene products. This changes during the maternal-to-zygotic transition (MZT) [18, 19], where the developmental control is gradually passed to the zygotic genome, while a large subset of maternal factors is cleared. Accordingly, the MZT is characterised by two phases: the maternal clearance and the zygotic genome activation (ZGA) [20].

The maternal transcripts cover a large fraction of the genome, about 70% in zebrafish embryos (ZFE), but this number varies across species [21]. The maternal clearance can be subdivided into two distinct pathways: an early maternally directed wave of decay and a later wave that requires zygotic products to promote the destabilisation of maternal transcripts [22]. In zebrafish, about two-thirds of cleared maternal transcripts degrade between the 1-cell and 16-cell stage (maternal pathway), while the remaining 40% are attributed to the zygotic pathway [21]. It is, however, likely that many maternal mRNAs undergo combined maternal and zygotic decay modes. Consequently, different groups of degradation profiles of maternal transcripts are observed in *D. rerio*: degradation shortly after fertilisation and degradation after ZGA. [23, 20, 24].

The onset of zygotic transcription does not occur until several hours post-fertilisation (hpf), with the timing varying across species [25]. Importantly, the ZGA is not a singular event, but rather a period, consisting of a minor and major transcriptional wave. The major wave of ZGA coincides with the mid-blastula transition after cell cycle 10, at approximately 3 hpf [19, 26]. The minor wave begins during early developmental stages and involves only a small subset of genes that are subsequently required for the full genome activation [20].

Considering these dynamic changes, stress regulator levels can vary significantly across early developmental stages, implying that the stress response is strongly dependent on exposure timing [27]. Assuming a constant baseline may therefore overestimate the detoxification capacity at stages where stress regulator concentrations are naturally low [10, 13]. Insights from dynamic RNA-based toxicokinetic-toxicodynamic (TKTD) modelling applied to *D. rerio* exposed to toxicants also suggested that accurate whole-organism transcriptome baselines are critical to separate toxic effects on transcriptome dynamics from developmental dynamics [28]. In this context, especially mechanistic baseline models (as opposed to purely empirical interpolations or splines) are superior representations as they provide the interface to describe feedbacks between intervention (e.g. toxicant exposure) and deviations from the transcriptional developmental norm.

Fortunately, baseline RNA transcriptomes of the early ZFE expression have recently become available [29, 30] and provide a valuable source to derive transcriptional development profiles for *D. rerio* during 120 hpf. Therefore, we propose a mechanistic transcription model, named *MZ-Baseline* model that captures common expression patterns and incorporates MZT dynamics to generate biologically informed baselines, describing the total transcript abundance as the sum of maternal and zygotic contributions.

This study aims to provide a biologically informed framework for understanding and modelling baseline gene expression dynamics in zebrafish during early development, thereby providing a foundation for integrating temporal baseline expression in toxicological research.

For this we fitted the *MZ-Baseline* model, as well as a commonly used single-state kinetic model for performance comparison, to a 120 hpf time-course RNA-seq dataset that combines measurements of over 30,000 genes from four independent studies. To compare the mechanistic approach with a shape-based one, we further we derived smooth expression trajectories from the dataset by fitting it to a generalized additive model (GAM). Using fuzzy *c*-means clustering (FCM), we identified common expression pattern in the developmental zebrafish transcriptome based on the shape similarity of the expression trajectories. We further applied FCM clustering to the mechanistic model parameters that describe the transcription and degradation rate as well as the temporal activation and regulation of each gene, to derive mechanistically informed expression cluster. These parameter-based clusters were then systematically compared to the shape-based approach.

## 2 Methods

Following the hypothesis that developmental genes of certain functional groups are expressed in distinct temporal pattern, we first applied fuzzy c-mean clustering (FCM) to identify common expression profiles during zebrafish embryogenesis. To first gain a general overview of common temporal pattern, we used smoothed, GAMs-fitted trajectories to identify expression clusters using FCM. Then, we fitted an mechanistic ODE-based model to identify parameters of temporal biological regulation as well as the maternal clearance mechanism for each gene. Subsequently, we used the fitted mechanistic model parameters to re-cluster the transcriptomics dataset using FCM.

### 2.1 Zebrafish culture

A wild-type adult zebrafish strain (*D. rerio*, strain OBI/WIK, generation F3), originating from an in-house cross of zebrafish strains UFZ-OBI and WIK, was used for all experiments. Fish were fed twice daily with dry food (SDS-400, Special Diets Services, Witham, Essex, UK) and once daily with *Artemia* sp. Fish were cultured under a 14 h light/10 h dark photoperiod in 20 L aquaria with a density of 1–2 fish per liter (*±* 26.5 1 °C). Water quality parameters were pH 7–8; water hardness 2–3 mmol/L, conductivity 540–560 *µ*S/cm, nitrate *<* 2.5 mg/L, nitrite *<* 0.025 mg/L, ammonia *<* 0.6 mg/L, oxygen saturation 87–91%. Spawning trays were inserted 4–6 h before the end of the light cycle. The following day, spawning was initiated by the onset of light and eggs were collected within 1 h. Facilities for breeding and the production of embryos were licensed by Landesdirektion Leipzig (Aktenzeichen 75-9185.64).

### 2.2 RNA extraction and sequencing

Embryos were incubated at 26 °C in glass vials with 10 embryos per vial. Two vials (20 zebrafish embryos in total) were harvested per replicate. Embryos were transferred into FastPrep tubes (MP Biomedicals, Irvine, CA, USA), media was removed, 150 *µ*L ice-cold TRIzol (Invitrogen, Thermo Fisher Scientific, Waltham, MA, USA) was added and samples were stored at −80 °C until further preparation. Homogenization was performed by a FastPrep-24 5G (MP Biomedicals, Irvine, CA, USA) for 30 s at 6.0 m/s two times. In between both steps the samples were incubated on ice for 1 min. The samples were centrifuged for 5 min at 4 °C at 12,000 *g*. Supernatant was transferred into a fresh 1.5 mL Eppendorf tube. RNA isolation was performed using a pipetting robot (Microlab STAR, Hamilton Company, Reno, NV, USA) following the manual provided for the MagMAX-96 for Microarrays Total RNA Isolation Kit (Invitrogen, Thermo Fisher Scientific, Waltham, MA, USA) and conducted in a 1.5 mL Eppendorf tube. RNA was eluted with a volume of 50 *µ*L. The quality of isolated RNA was assessed using a NanoDrop spectrophotometer (Thermo Fisher Scientific, Waltham, MA, USA) and a 2100 Bioanalyzer (Agilent Technologies, Santa Clara, CA, USA) with the Agilent RNA 6000 Nano Kit. Samples with an RNA integrity number (RIN) greater than 7 and calculated concentrations exceeding 25 ng/*µ*L were used for further processing. All samples were diluted to a concentration of 25 ng/*µ*L (10 *µ*L in total) by adding RNase-free water. Poly(A)^+^ mRNA was enriched using the NEBNext Poly(A) mRNA Magnetic Isolation Module (New England Biolabs, Ipswich, MA, USA). Sequencing libraries were generated using the NEBNext Ultra II Directional RNA Library Prep Kit for Illumina (New England Biolabs, Ipswich, MA, USA), and index codes were added to attribute sequences to each sample. The library preparations were sequenced using the Illumina NovaSeq 6000 platform (Illumina Inc., San Diego, CA, USA) with a paired-end sequencing length of 100 bp (PE100). The sequencing was performed at ParoX GmbH (Leipzig, Germany). Datasets BK and JN are made publicly available at GEO with accessions GSE352672 and GSE352674, respectively.

### 2.3 Bioinformatic data processing

All datasets used in this study were processed from raw FASTQ files. Adapter sequences were trimmed from raw reads with fastp (v0.24.0). Transcript abundances were quantified with Salmon (v1.10.1) against the Ensembl zebrafish transcriptome (release 111) based on the GRCz11 assembly, using an index built with a k-mer size of 31.

### 2.4 Dataset

We used an time-resolved bulk RNA-sequencing dataset of zebrafish developmental stages from 0 to 120 hpf. The dataset comprises expression data of 36,570 genes from 4 sources (White et al., Pauli et al, BK, JN) with different number of samples and sampling times each, adding up to 50 unique time points (Tab. 1). Transcript-level measurements were aggregated to gene level by summing the expression levels (TPM, transcripts per million) of all transcripts associated with the same gene. Furthermore, replicate samples were aggregated per source by taking the arithmetic mean of gene-level TPM values.

Developmental rate of zebrafish embryos increases with higher incubation temperature. Nominal sampling times under different incubation temperatures therefore describe different developmental stages. To address this, sampling times were converted to standard developmental time using the temperature-corrected formula by Kimmel et al.[1]:

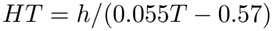

with HT = hours of development at temperature T and h = hours of development to reach the stage at 28.5 °C. The four datasets showed overall good alignment as PCA analysis shows (Fig. 1), with the developmental time explaining 64.8% of variance in the dataset. Only three clear outliers were identified (BK 5.16 hpf, BK 8.6 hpf and Pauli et al. 120 hpf), which were kept in the joint dataset.

**Figure 1.**
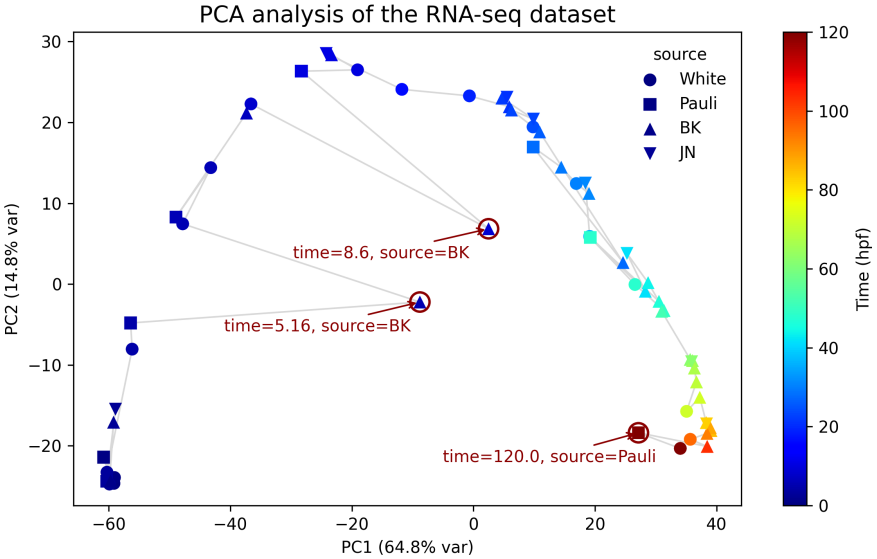
PCA analysis of the combined dataset with source and time samples. With the exception of three outliers (BK 5.16 hpf, BK 8.6 hpf and Pauli 120 hpf) the four datasets are well aligned, showing similar expression profiles across the zebrafish developmental time course. PC1 captures the developmental time, explaining 64.8% of the variance in the data.

**Figure 2.**
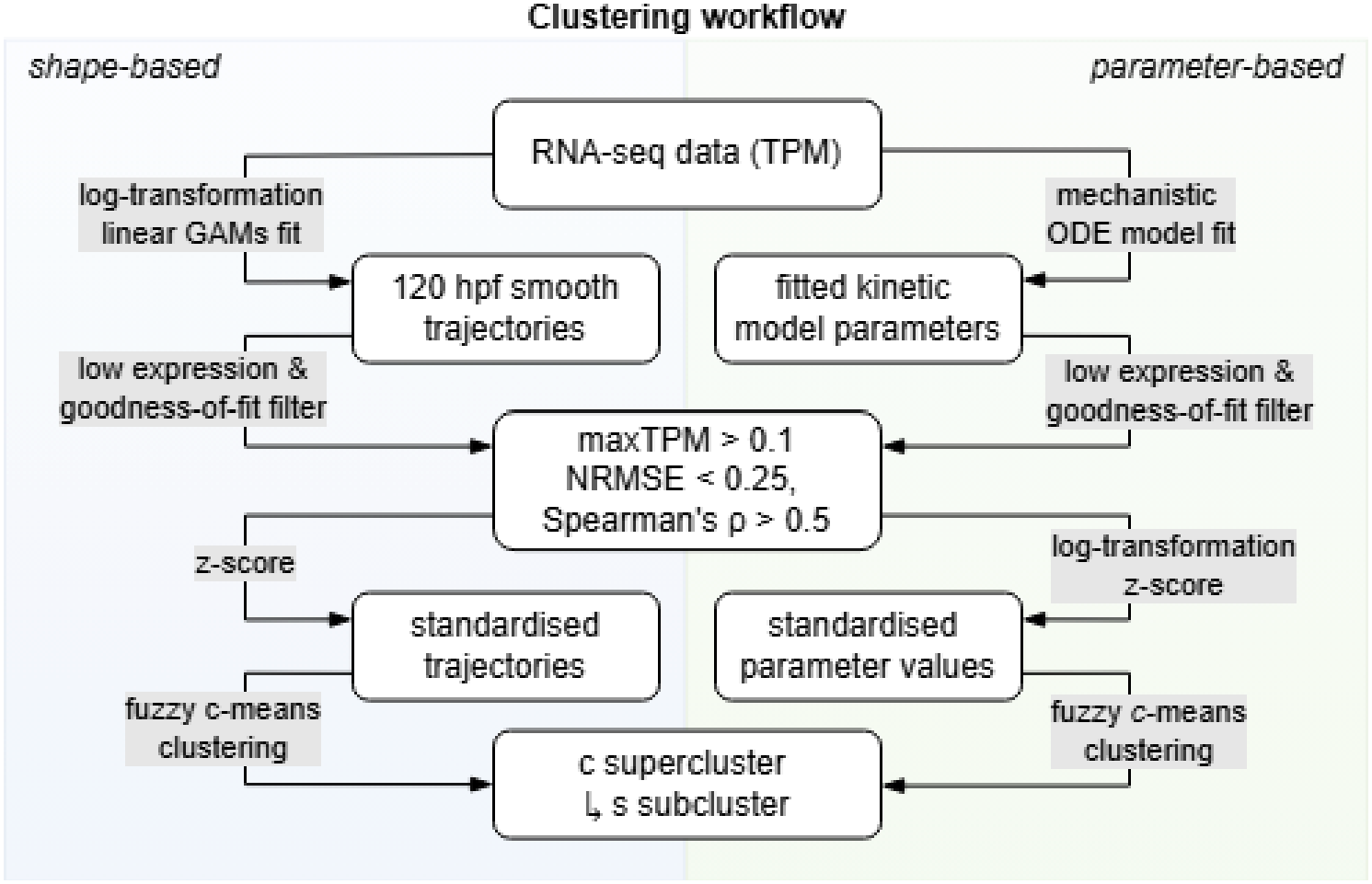
Computational workflow for clustering of gene expression baselines using FCM.

**Table 1.** The joint dataset is composed of four sources with different sample resolutions of ZFE stages between 0 and 120 hpf.

| Source | Samples | Time span<br>(hpf) | Replicates |
|---|---|---|---|
| White et al. | 18 | 0 - 120 | 5 |
| Pauli et al. | 8 | 0.875 - 120 | 8 |
| BK | 28 | 1.29 - 103.2 | 1 - 7 |
| JN | 8 | 2.58 - 82.56 | 2 - 10 |

### 2.5 Fuzzy c-means clustering

To identify common expression profiles of developmental genes and compare a shaped-based versus a parameter-driven approach, we applied fuzzy c-means clustering (FCM) [31] both on expression trajectories and on fitted *MZ-Baseline* model parameters. Since gene expression data is noisy and the expression levels can differ in order of magnitudes across genes, we established a workflow for harmonising and clustering the dataset (see 2).

As a first step in the shape-based approach, we log-transformed the data and fitted a Generalised Additive Model (GAM) to smoothen the gene expression trajectories prior to clustering and generate harmonised sampling times. We used the implementation of the Python package *pyGAM.LinearGAM* [32] in the form of:

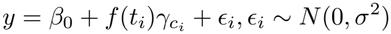

Here, *β*_0_ is an intercept, *f* (*t*_*i*_) is a smooth function of time, expressed as a sum of *K* b-spline basis functions shared across sources, 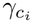is a factor effect adding a separate offset for each source *c*_*i*_, and *ϵ* is Gaussian observation noise. The number of b-spline basis functions *K* was limited to 20 splines to prevent overfitting.

For the parameter-based approach, a mechanistic ODE model was fitted to the untransformed TPM data. Then, in the second step, the model fit quality was assessed. Because noisy, low expression data can distort the result of the clustering analysis, we applied a low-expression filter to remove all genes whose maximum expression is below 0.1 TPM in any source. This reduced the dataset to a total of 29,308 genes. Furthermore, we assessed the goodness-of-fit using the NRMSE and Spearman’s *ρ*. Model fits that did not pass the goodness-of-fit criteria were excluded from the FCM clustering analysis to prevent distortion of the clusters by unreliable, poor-fitted data.

In the third step, the expression trajectories and fitted model parameters were standardised by applying z-scoring (*Z* = (*x−µ*)*/σ*). This enables the clustering algorithm to focus on the shape of the expression profiles rather than expression magnitude.

Fuzzy *c*-means (FCM) is an unsupervised clustering algorithm that partitions the dataset into *c* clusters by assigning each gene a cluster membership [31]. It is described by the following objective function:

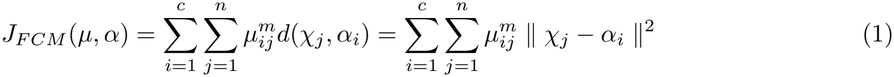

with the weighting index *m >* 1 controlling the fuzziness, *µ*_*ij*_ being a fuzzy *c*-partition and *α* being a set of *c* cluster centers. Fuzzy *c*-means was applied using the python package *scikit-fuzzy* [33].

The cluster with the highest membership degree was hard labelled as the genes cluster. As the number of clusters *c* is unknown for the given dataset, we used the Xie-and-Beni index (XB-index) to determine the optimal number of clusters. The XB-index is a validity measure for fuzzy clustering methods that compares the “compactness” *J*_*m*_ (distance within the cluster) and the “separation” (*d*_*min*_)^2^ (distance between clusters) for each of *n* data vectors, using euclidean distance measures. It is defined as

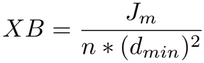

where the smallest XB value indicates the optimal number of clusters [34]. To find the optimum, we applied FCM clustering over a range of c=2 to c=9 clusters, selecting the *c* yielding the lowest XB-value for final clustering. For validation of the choice, we computed the cluster stability [35]. This method estimates the clustering reproducibility for two random subsamples of genes using bootstrap resampling for a number of clusters *c*. The cluster centers are projected onto the full dataset, and the assigned cluster label is compared with the Adjusted Rand Index (ARI). A higher ARI means higher consistency across subsample clustering, meaning higher stability of cluster assignment. The ARI was implemented using the *adjusted rand score* from *sklearn.metrics* [36]. Furthermore, we reported the mean fuzzy partition coefficient (FPC), which indicates by values between 0 and 1 how clear the cluster assignment is. A FPC values close to 1 corresponds to distinct groups. For definition of subclusters, we re-applied FCM clustering for each supercluster partition to identify underlying expression pattern. Next to the number of cluster *c*, the fuzziness exponent *m* is another important parameter in FCM. For this analysis, *m* was set to 1.6.

### 2.6 Baseline transcription model

A commonly used mathematical model to describe changes in transcript concentration integrates the basic kinetics of transcription and RNA degradation [37, 38, 39]. Transcription is assumed to be independent of the current RNA abundance and is therefore modelled as a zero-order process with rate *β*, while RNA decay follows first-order kinetics with constant rate *δ*. This leads to the following ODE for changes in mRNA concentration:

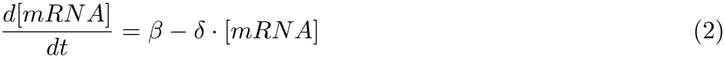

While the *Basic* model captures the basal gene expression kinetics, it does not account for the specific and complex dynamics of the MZT. During the early stages of embryogenesis, maternal factors drive the developmental processes and thus determine the initial transcript levels. In zebrafish, maternal transcripts cover approximately 70% of the transcriptome [21]. During maternal clearance, a subset of these maternal factors is eliminated. This event can be subdivided into two distinct pathways: an early, maternally directed wave of decay and a later wave that requires zygotic products to promote the destabilisation of maternal transcripts [22]. In zebrafish, about two-thirds of cleared maternal transcripts degrade between the 1-cell and 16-cell stage (maternal pathway, M-decay), while the remaining 40% are attributed to the zygotic pathway (Z-decay) [21]. The maternal pathway is regulated, among others, by RNA-binding proteins (RBPs). In the oocyte, RBPs stabilise and repress maternal mRNAs to prevent their translation, while during embryogenesis, they can coordinate their degradation [40, 16]. One of the principal promoters of the zygotic pathway are miRNAs, e.g. miR-430 as one of the first zygotic transcripts in zebrafish. miR-430 binds specific motifs in the 3’ UTRs of maternal mRNAs, which enables their selective targeting. Mechanistically, miR-430 binding first causes translational repression and then recruits the CCR4–NOT deadenylase complex, leading to poly(A)-tail shortening, destabilisation, and subsequent degradation of the targeted transcript [41, 42]. Furthermore, AU-rich element binding proteins (ARE-BPs) have been identified to play a role during maternal clearance in zebrafish [16, 21].

Thus, following the described theory, maternal transcripts with zygotic degradation pathway are expected to remain at a steady level before ZGA, while transcripts following the maternal degradation pathway are degraded shortly after fertilisation. As they are deposited during oogenesis and not transcribed de novo after fertilisation, their overall abundance decreases over time until complete depletion.

Following the activation of the embryonic genome a few hours post-fertilisation, zygotic mRNAs are synthesised and progressively dominate the expression dynamics [20, 21]. Consequently, the total mRNA pool transitions from purely maternal to a mixture of maternal and zygotic transcripts, and eventually to predominantly zygotic transcripts.

To take these dynamics into account, a two-state ODE model is proposed, hereafter referred to as **MZ-Baseline** model, with variants *M-decay* and *Z-decay*. Maternal transcripts, denoted by [*M*], are modelled using first-order degradation kinetics. In the *M-decay* model, the degradation rate *δ*_*M*_ is assumed to be constant, whereas in the *Z-decay* model, degradation depends on the activity of a zygotic regulator *R*(*t*).

For zygotic transcripts, transcriptional activation occurs at *t*_*zga*_, following a zero-order kinetic with rate *α* [43]. The activation process is modelled as a logistic function *σ*, which is defined by parameters *t*_*zga*_ for the temporal onset of transcription and the steepness *s*. At time *t*_*reg*_, the transcription factor is assumed to reach a critical threshold concentration, resulting in a change of the transcription rate to *β*, representing either repression (*β* ¡*α*) or activation (*β* ¿ *α*). For the special case *β* = *α*, the transcription rate does not react to the transcription factor but remains at the initial rate.

Degradation of zygotic transcripts follows first-order decay at rate *δ*_*Z*_. The total mRNA concentration of a gene is computed as the sum of maternal and zygotic transcripts (*y*(*t*) = *M* (*t*) + *Z*(*t*)).

The *MZ-Baseline* model is described by the following equations:

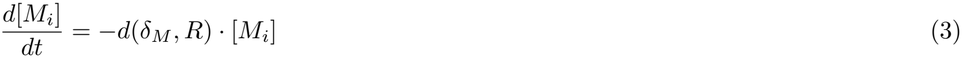

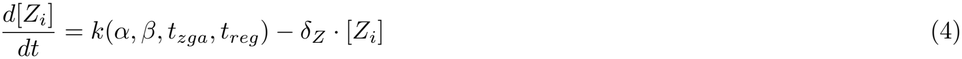

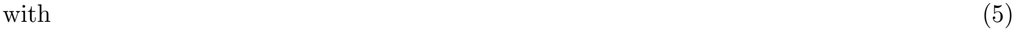

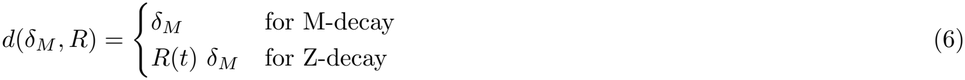

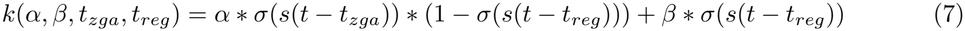

Figure 3 show the exemplary model outcome for the two variants M-decay and Z-decay and for three different regulatory modes (upregulation, downregulation, and no regulation) for maternal and zygotic transcripts, respectively.

**Figure 3.**
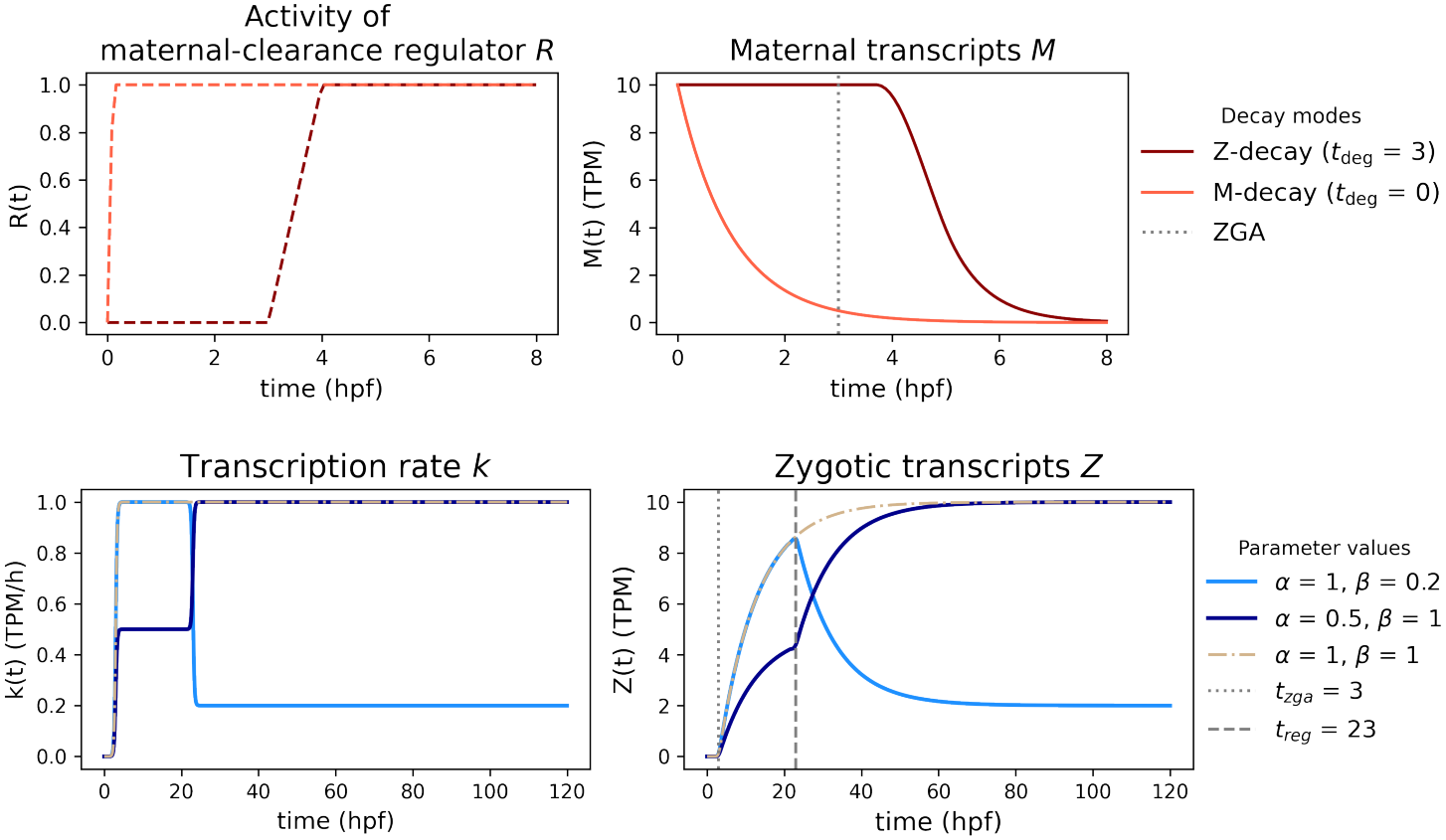
Upper Figure: Maternal transcripts are targeted for degradation via either the maternal (M) or zygotic (Z) decay mode. For the M-decay model, the maternal repressor is assumed to be present from 0 hpf, while the zygotic repressor is accumulated and activated during ZGA at 3 hpf (left Figure). This produces either immediate or delayed degradation of maternal transcripts (right figure). Lower Figure: Transcription rate of the *Baseline* model. The dark blue line represents transcriptional downregulation (*α > β*), the light blue line represents transcriptional upregulation (*α < β*) at *t*_*reg*_. The dashed line represent the special case of no regulation (*α* = *β*).

### 2.7 Parameter estimation and model implementation

A key challenge in fitting large-scale transcriptomic datasets is the wide range in transcript abundances, which can span several orders of magnitude. Consequently, priors must be adjusted individually to gene-specific, plausible ranges that reflect the expected expression levels.

For the *MZ-Baseline* model, the initial maternal transcript abundance *M*_0_ is fixed to the observed initial value, whereas the initial zygotic transcript level *Z*_0_ is assumed to be zero.

Turnover rates of individual mRNAs can vary substantially, with half-lives ranging from several minutes to multiple hours depending on transcript structure and sequence, the cellular environment or function [44, 45, 46]. In general, the degradation rate can be approximated from the mRNA half-life *t*_1*/*2_, via *δ* = *ln*(2)*/t*_1*/*2_ [38]. For M-decay transcripts, maternal mRNA is assumed to be primarily degraded prior to ZGA, with an estimated half-life of 2 h. This corresponds to a degradation rate of *δ*_*M*_ = *ln*(2)*/*5*h ≈* 0.35*h*^−1^. Accordingly, the prior for the maternal degradation rate in M-decay models is defined as a log-normal distribution with a mean of 0.35 and a standard deviation of *σ* = 1.

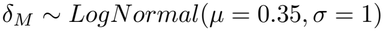

During MZT, maternal transcripts are actively targeted by miR-430, leading to accelerated decay and shorter half-lives. miR-430-mediated targeting begins at approximately 4 hpf, achieving complete deadenylation by around 6 hpf [41]. For Z-decay models, the half-life of targeted mRNAs is estimated to be 0.5 h. The maternal degradation rate is thus initialized with *δ*_*M*_ = *ln*(2)*/*0.5 *h≈* 1.4*h*^−1^. The degradation rate of zygotic transcripts, *δ*_*Z*_, is fixed at 0.1 h^-1^, corresponding to high stability with an assumed half-life of 7 h.

For initialisation of the transcription rate *β*, the steady state solution of the *Basic* model is used: *Z*^*∗*^ = *β/δ*_*Z*_. For each gene, *Z*^*∗*^ is computed as the mean observed expression level of the final three time points, multiplied by the initial degradation rate to obtain *β*. For the *MZ-Baseline* model, the initial transcription rate *α* is scaled to the maximum observed expression *Z*_*max*_ in the interval t = 2.25 to 72 hpf.

This results in the following priors:

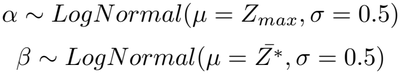

The timing parameter *t*_*reg*_ is defined as *t*_*zga*_ + *dt*_*reg*_ to ensure *t*_*zga*_ *< t*_*reg*_. For prior definition, the ZGA onset *t*_*zga*_ is centred at 3 hpf, while the regulation onset *t*_*reg*_ is initialised at later timing:

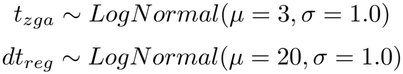

The steepness parameter *s* of the sigmoid activation function *σ*(*x*) is fixed at a value of 4, which relates to an activation window of approximately 1 h. For all models, we used a normal distribution as error model with *σ*_*y*_ ~ *LogNormal* as scale, and assumed a multiplicative error, defined as

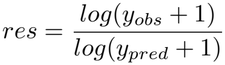

An overview of the model parameters can be found in S1.

For the evaluation of different aspects of the fit quality and for model selection, we choose a set of goodness-of-fit metrics 2. For convergence diagnostics, we chose 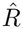 and the effective sample size (ESS). As model selection criteria, that balances model complexity and fit quality, we chose the Bayesian Information Criterion (BIC). To assess the quantitative model error, the Normalised Root Mean Squared Error (NRMSE) was used and for assessment of the qualitative pattern accuracy, we chose the Spearman’s *ρ* rank coefficient. A detailed explanation of the metrics can be found in S2.

The models were implemented in Python, using the model-building platform pymob [47]. Pymob provides several inference backends, including numpyro, which supports Bayesian inference algorithms such as Markov Chain Monte Carlo (MCMC) and stochastic variational inference (SVI). Parameter estimation was conducted using the integrated MCMC sampler NUTS with 4 parallel chains. The model fits were performed on the HPC of Osnabrück University.

## 3 Results

### 3.1 Shape-based FCM clustering reveals three expression clusters

Of the GAMs fitted trajectories, 29,144 genes in total passed the goodness-of-fit evaluation (Fig. S1) and were used in the clustering analysis. Fuzzy *c*-means (FCM) clustering identified three expression supercluster (SD, SU and DSU), which we named according to their overall observed expression pattern.

Figure 4 shows the trajectories of each supercluster center alongside with the average of the corresponding observed data.

**Figure 4.**
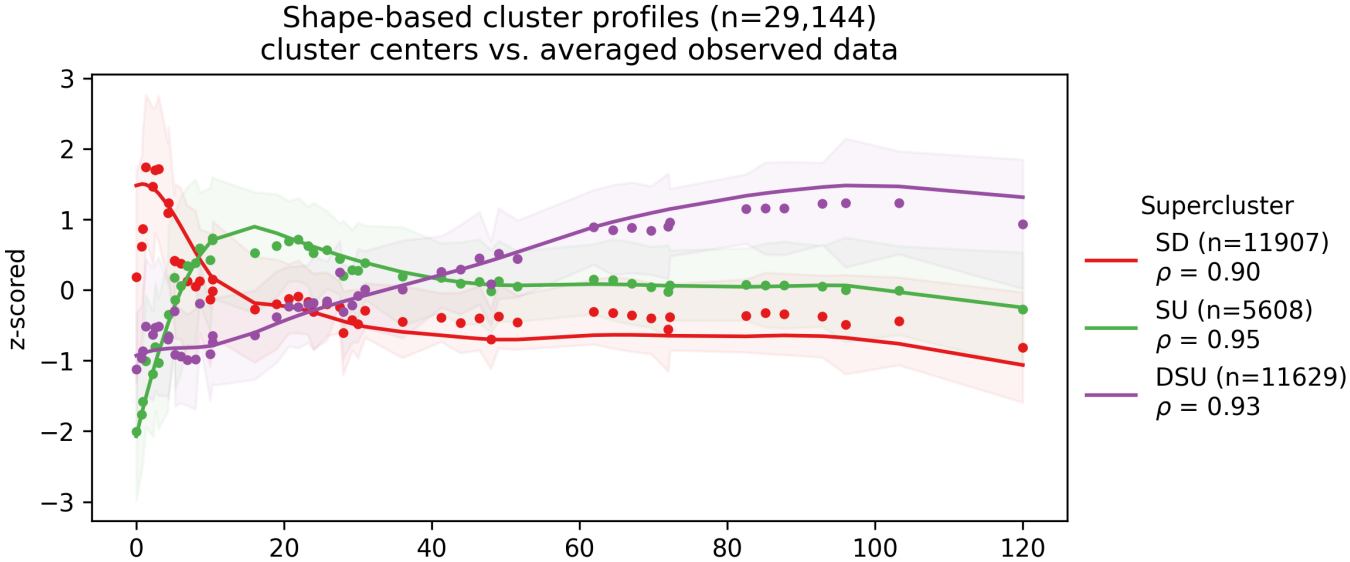
The Figure shows the supercluster centers (lines) and the average observed expression (dots) per supercluster with Spearman’s *ρ* for pattern accuracy. The bands show the standard deviation of observed expression per cluster. Expression levels are log2(TPM+1) scaled and standardised by z-score.

The **sustained downregulation (SD)** cluster is the largest cluster, comprising 11,907 genes (40.9% of the analysed dataset). The cluster center’s trajectory starts at high initial expression and decreases immediately to low levels that plateau around 20 hpf. The SD cluster center shows a high positive pattern correlation (Spearman’s *ρ* = 0.9) with the observed average. Despite this, the supercluster center appears to underestimate early dynamics, where the observed averages visibly shows a transient increase. Subclustering revealed two SD subclusters (see Fig. S3) with distinct dynamics. Genes in subcluster SD-1 (n=6,413) decrease steadily towards zero after a slight early increase, whereas genes in subcluster SD-0 (n=5,494) are transcribed following an initial decrease, stabilizing at levels below the initial expression level.

The **sustained upregulation (SU)** cluster center (5,608 genes; 19.2%) starts at low expression levels and increases immediately, reaching a plateau at around 20 hpf. It shows a high pattern accuracy (*ρ* = 0.95) compared to the observed average expression, which can also visibly be confirmed. Subclustering revealed two SU subclusters that differ in both transcription rate and long-term stability. The subcluster center of SU-0 (n = 2,460) rises more quickly but subsequently decreases, stabilising below its early peak, while SU-1 (n = 3,148) increases more slowly and remains comparatively stable thereafter.

The **delayed sustained upregulation (DSU)** cluster (n = 11,629 genes; 39.9%) starts at low expression levels, and only shows significant increases at around 10 hpf, which is several hours after the canonical timing of zygotic genome activation (ZGA) in zebrafish at 3 hpf. The supercluster center reaches its plateau around 100 hpf. Overall, the DSU cluster center shows high correlation with the observed cluster average (*ρ* = 0.93), although early expression dynamics until approximately 10 hpf appear to remain uncaptured. Subclustering identified three DSU subclusters, that differ significantly in early dynamics. DSU-0 (n = 4,556) and DSU-2 (n = 5,303) both start at low expression levels and increase thereafter, with DSU-0 genes increasing more rapidly than DSU-2 genes. Compared to that, DSU-1 (n = 1,770) genes start at higher expression, decrease initially, and then increase again towards a high, stable plateau.

Overall, the three superclusters showed a high mean internal FCM membership (SD 88%, SU 76% and DSU 86%, see Fig. S4), confirming that there is little overlap with other clusters and most genes can be distinctly attributed to one characteristic expression profile. Despite this, the membership degree revealed a slight overlap of the SU and SD cluster (15% and 10%), indicating that some genes likely show a more complex pattern that cannot attributed to one dominant cluster only.

### 3.2 *MZ-Baseline* model outperforms the *Basic* kinetic model - model evaluation and selection

The mechanistic *MZ-Baseline* and the kinetic *Basic* model were fitted and evaluated on the full non-zero dataset (n = 31,658 genes). The model fits were evaluated against convergence criteria 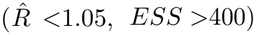 and fit-quality criteria, considering a quantitative (*NRMSE <*0.3) and qualitative (*ρ >*0.5) goodness-of-fit threshold. A fit was flagged as “fully accepted” when all criteria were met. The *MZ-Baseline* model yielded fully accepted fits for 55,3% (n = 18,013) of genes with the Z-decay variant and 55.0% (n = 17,923) with the M-decay variant. Under the *Basic* model approach (zero-order production, first-order degradation) only 23.6% (n = 7,814) of model fits met all goodness-of-fit criteria (Fig. S6). The *MZ-Baseline* model showed more convergence failures than the *Basic* model, but far fewer poor-quality fits: 15.1% of M-decay and 13.3% of Z-decay fits were rejected for poor fit quality, compared to 37.5% for the *Basic* model. As with the GAM fits, ODE model fits were accepted if they passed either the quantitative or the qualitative criterion.

Fit quality varied among expression superclusters (Fig. 5). The *Basic* model performed particularly poorly with pattern reproduction in the SD and SU clusters, with acceptance rates of only 34.19% and 20.93% respectively. Both *MZ-Baseline* model variants achieved overall good pattern accuracy in the SU and DSU clusters, with acceptance rates between 82% and 87%. In the SD cluster, only 65.2% and 68.5% of the M-decay and Z-decay variant, respectively, are able to reproduce the pattern. Considering quantitative fit quality alone, however, the SD cluster achieved the highest acceptance rate for all model variants.

**Figure 5.**
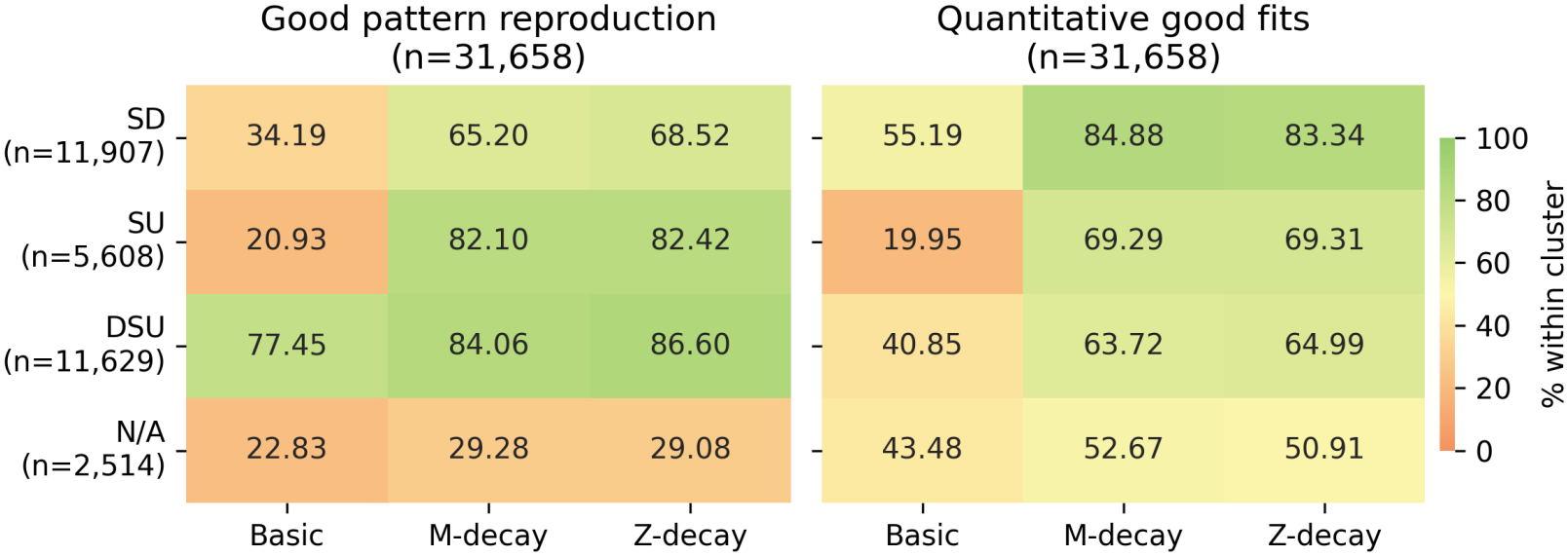
Left Figure: Baseline model fits producing good expression pattern reproduction, assessed by Spearman’s *ρ >* 0.5. Right Figure: Baseline model fits producing quantitative good fit with low residual error, assessed by NRMSE *<* 0.25. The N/A block comprises the genes, that were excluded from the clustering analysis due to poor fit or expression levels below minimum threshold.

**Figure 6.**
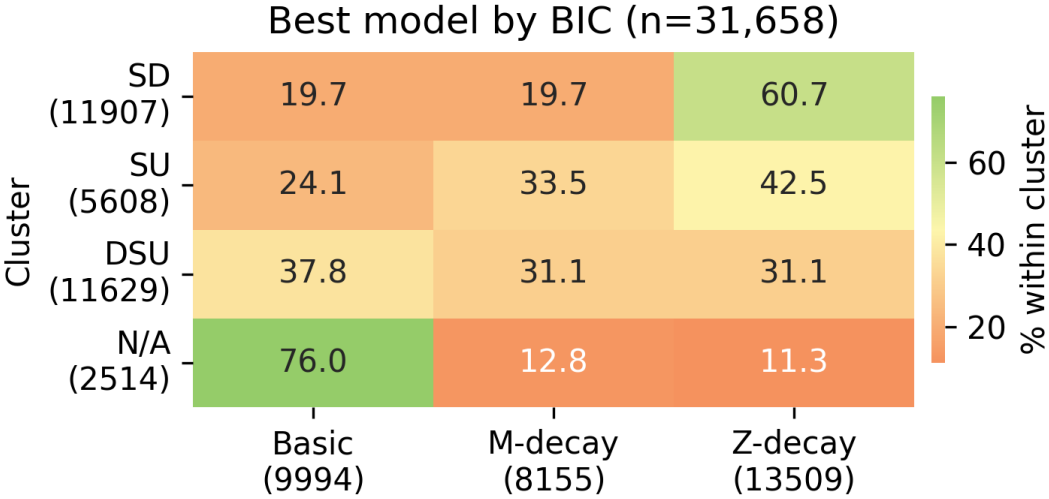
Model with lowest BIC was chosen as best fitting model. Despite the higher model complexity (5 free parameter), the *MZ-Baseline* model achieved a high preference rate compared to the simpler *Basic* model (2 free parameter). The N/A block comprises the genes, that were excluded from the clustering analysis due to poor fit or expression levels below minimum threshold.

We used the *BIC* to select the best-fitting model for each gene (Fig. **??**). The*Z-decay* model variant was preferred for 42.7% of genes, of which 51.7% showed strong evidence over the *Basic* model (Table 3). The *M-decay* variant was preferred for 25.8% of genes, of which 58.9% showed strong evidence. For the remaining 31.6% of genes, the simpler *Basic* model was able to sufficiently represent the expression pattern.

**Table 2.** Goodness of fit metrics used for evaluation of Baseline model fits.

| Metric | Used for | Threshold | Equation |
|---|---|---|---|
| R-hat ($\hat{R}$) | Convergence diagnostic | < 1.05 | arviz_stats.rhat |
| ESS (Effective Sample Size) | Convergence diagnostic | > 400 | arviz_stats.ess |
| PPC (Posterior predictive coverage) |  |  |  |
| BIC (Bayesian Information Criterion) | model selection | ΔBIC>6 | $-2ln(\hat{L} + k * ln(n))$ |
| NRMSE (Normalised Root Mean Squared Error) | quantitative fit accuracy | < 0.3 | $\frac{1}{y_{\max} - y_{\min}}\sqrt{\frac{\sum_{i=1}^{n}(y_i - \hat{y}_i)^2}{n}}$ |
| Spearman's ρ | qualitative shape accuracy | > 0.5 | scipy.stats.spearmanr |

**Table 3.** BIC report. Transcriptome-wide median BIC values and 90% percentile interval (pi) (lower 5%, upper 95%) per model are reported for a 120 hpf time course with 50 observation points and 31,658 genes. *k* denotes the number of free model parameters. Δ*BIC*_*i*_ *>* 6 refers to comparison to the *Basic* model, thus the percentage of genes showing strong evidence for the *MZ-Baseline* model.

| Model | k | Median BIC | 90% PI | preferred gene-wise | Δ*BIC*<sub>*i*</sub> > 6 |
|---|---|---|---|---|---|
| Basic | 2 | 76.91 | -175.62 181.63 | 31.56% | - |
| Baseline-M | 5 | 53.24 | -172.29 157.74 | 25.75% | 58.90% |
| Baseline-Z | 5 | 49.64 | -171.35 154.46 | 42.67% | 51.68% |

Model preference also depended on the expression cluster and abundance. The *Basic* model was preferred almost exclusively for low-expressed genes excluded from the shape-based clustering analysis (76% preference within this group), and remained competitive in the DSU cluster, where it was preferred for 38% of genes under consideration of model complexity. This indicates that a third of genes follow likely a rather simple upregulation pattern, which can sufficiently be described through basic kinetics. In contrast, the Z-decay variant dominated in the SD cluster, preferred for 61% of genes, potentially reflecting a genuinely dominant zygotic clearance mechanism among maternally deposited transcripts with high initial expression in this cluster.

Based on these results, all subsequent analyses focus on the accepted *MZ-Baseline* model fits, given their higher explanatory power for the majority of genes.

Apart from overall fit quality, the *MZ-Baseline* model allows us to infer the underlying maternal decay mechanism. For this, we used gene-wise comparison of the M-decay and Z-decay variants by BIC.

The acting maternal-clearance mode was identified using Δ*BIC >* 6 as an indicator of strong model evidence, comparing BIC values of M-decay and Z-decay model variants. Under this assumption, strong model evidence towards one of the decay modes was found in 35.3% of genes. Overall, 62.7% of genes showed a preference for the zygotic pathway, while the remaining 37.3% are best explained with the maternally directed degradation pathway. This is a higher proportion of Z-decay genes than reported by Vastenhouw et al. [21] (approx. 40%), suggesting that the *MZ-Baseline* model may overestimate the prevalence of the zygotically driven clearance mechanism.

This decay-mode preference varied by expression clusters (Fig. 7). Genes from the SD cluster indicate a clear preference for Z-decay (n = 7,934 vs. n = 3,112 for M-decay), with a high percentage of strong model evidence (64.9%). Because SD cluster genes are characterised by high initial expression levels, following degradation, the abundance of SD cluster genes during early developmental stages is predominantly composed of maternal transcripts. This benefits the model selection as the maternal dynamics are not overlaid with zygotic transcription.

**Figure 7.**
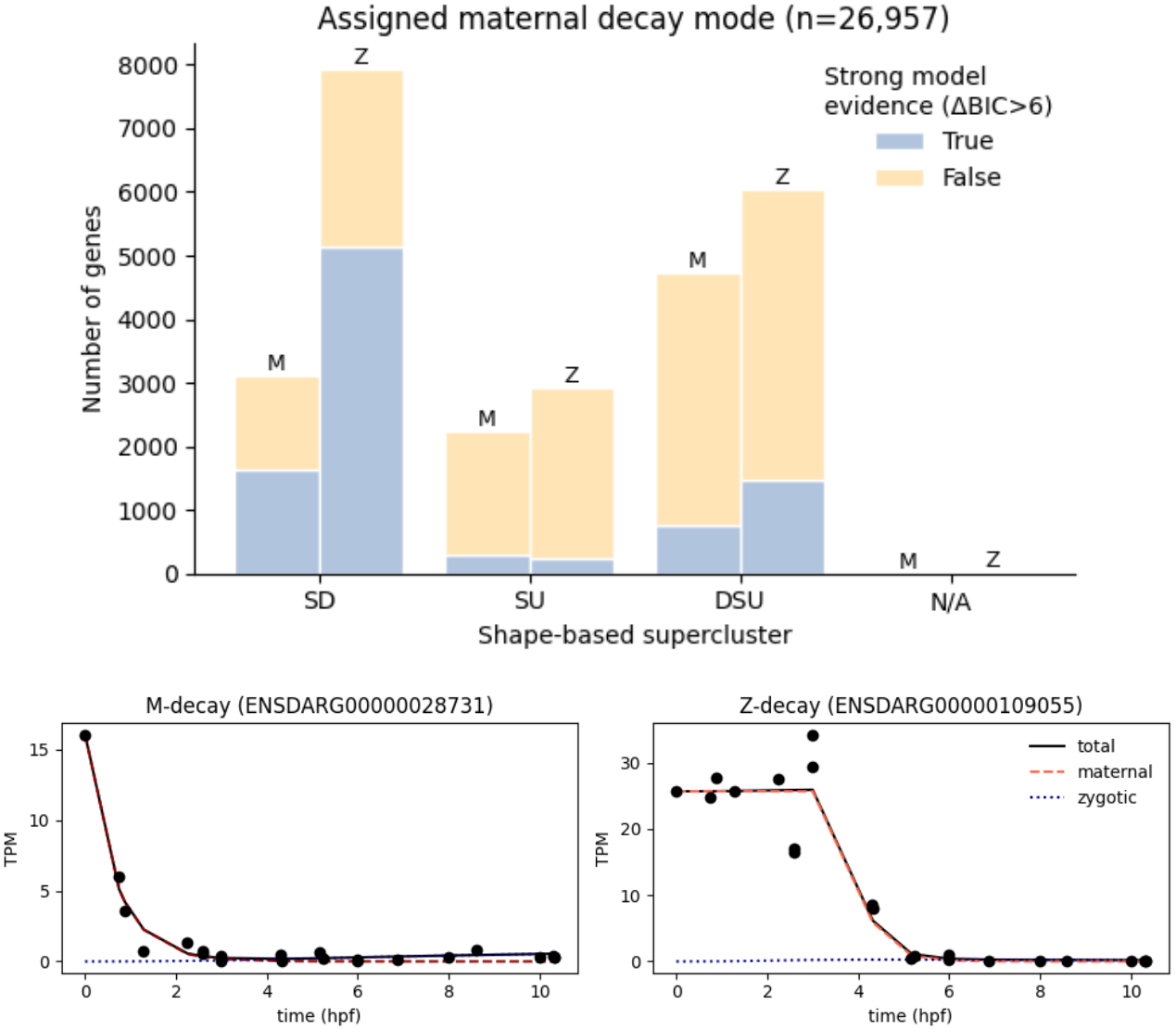
The upper figure shows the assigned maternal decay mode. The model variant with the smaller BIC value provides evidence for the acting maternal decay mode (maternal or zygotic repressor). A value of Δ*BIC >* 6 was interpreted as strong model evidence for the selected decay model. The lower figure shows the expression profile of two specific genes, over an 12 hpf time course with each the highest model evidence for the M- and Z-decay mode.

The DSU cluster showed a weaker preference for the Z-decay variant (n = 4,715 M-decay vs. n = 6,040 Z-decay), whereas the SU cluster shows no dominant preference for either mode, consistent with the predominantly weak model evidence in this supercluster. This is expected, as both SU and DSU genes typically contain a low to negligible abundance of maternally provided transcripts (with exception of the subcluster DSU-1), so maternal transcript are either largely absent or quickly replaced by zygotic transcripts, making it difficult to clearly identify the maternal decay dynamics.

Figure 7 illustrates the model selection with the expression profiles of two genes with each showing high model evidence towards one of the decay modes. Both follow the expected dynamics: Starting at initially high expression level, the M-decay gene (assigned to the SU supercluster) decreases immediately after fertilisation, reaching complete depletion by 3 hpf. The Z-decay gene (assigned to the SD cluster) on the other hand remains at relatively stable level until the onset of ZGA by 3 hpf.

Overall, these results demonstrate one of the mechanistic advantages of the *MZ-Baseline* model: unlike purely descriptive approaches, it allows the underlying maternal clearance mechanism to be inferred directly from single gene expression dynamics. For all subsequent analyses, each gene was assigned to either the maternal (M-decay) or zygotic (Z-decay) degradation pathway, based on the BIC-preferred model variant.

### 3.3 Mechanistic model parameter-based FCM clustering

For the parameter-based FCM clustering, genes that were flagged as poorly fitted in the previous evaluation as well as genes with a maximum expression below 0.1 TPM (as in the shape-based clustering) were excluded from the analysis to prevent distortion due to badly represented or too noisy baselines. This left a total of 26,957 genes that were included in the clustering analysis.

For parameter-based FCM clustering, each gene was represented by a set of inferred parameter from the *MZ-Baseline* model fits, namely the regulation indicator *r* = *β/α*, the maternal half-life *t*_*m,half*_ = *ln*(2)*/δ*_*m*_, the onset of zygotic transcription *t*_*zga*_, the regulatory transition time *t*_*reg*_, the time of peak expression *t*_*peak*_ and the initial expression level *y*_0_. The parameter values were log-transformed and z-scored prior to conducting FCM clustering. The distribution of the input parameters can be found in S7.

Four expression clusters were identified from the inferred *MZ-Baseline* model parameters. The supercluster centers of each parameter distribution as well as the mean model trajectory per supercluster are shown in Figure 8. Three of these clusters correspond to patterns already identified in the shape-based approach: sustained downregulation (SD), sustained upregulation (SU), and delayed sustained upregulation (DSU). The fourth is a newly resolved **transient upregulation (TU)** cluster, comprising genes that start at low expression, increase sharply immediately after fertilisation, peak around 4 hpf, and then decrease steadily.

**Figure 8.**
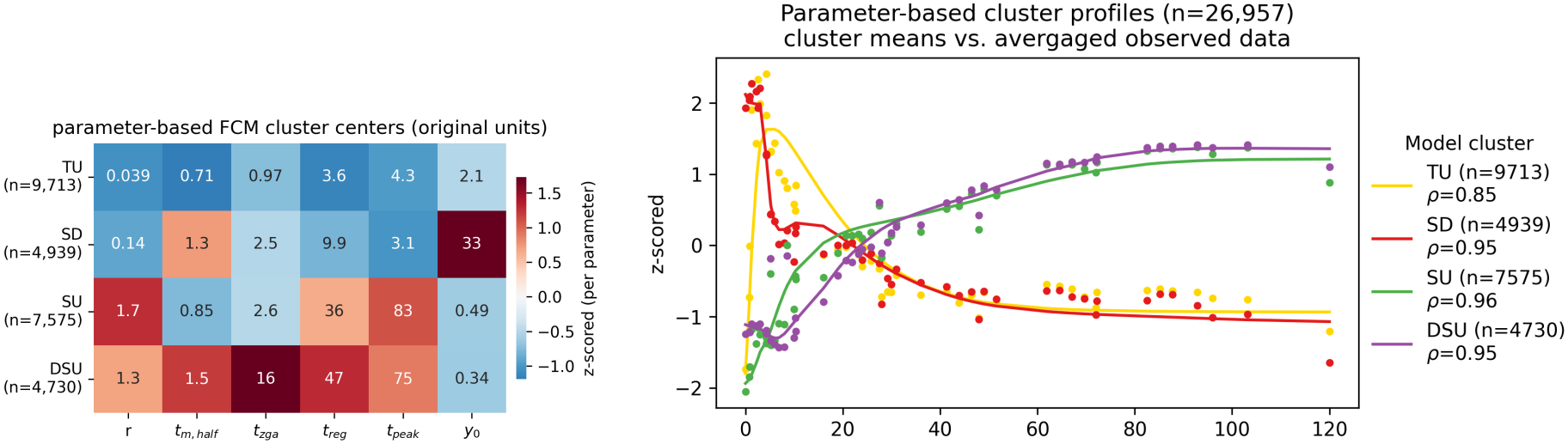
Left figure: Cluster centers of model parameters used in FCM clustering. The regulation indicator *r* is defined by the relation *β/α*. A value of *r >* 1 indicates upregulation at *t*_*reg*_ while a value of *r <* 1 indicated downregulation. Right Figure: Average cluster profiles of fitted Baseline-model trajectories (lines) and mean of observed data per supercluster (dots).

The supercluster means correlated closely with the observed average (Spearman’s *ρ* = 0.85 - 0.96), indicating overall high pattern accuracy. Notably, early dynamics were resolved substantially better than in the shape-based approach: The TU cluster captures the transient pre-ZGA increases, which the shape-based approach failed to represent. The transient decrease observed in the DSU cluster is likewise reflected in the supercluster centroid. Consistent with this, the mean of observed expression in the SD supercluster no longer shows the transient increase, which is also reflected in a higher pattern accuracy (increased *ρ*, from 0.9 in shape-based to 0.95 in parameter-based SD cluster).

The parameter-based cluster centers further highlight the mechanistic differences between clusters. The regulation indicator *r* is consistent with the overall observed pattern. Here, an *r* value greater than 1 indicates upregulation of the expression level at *t*_*reg*_ while a value of *r <* 1 indicated downregulation. The TU and SD cluster both report values of *r <* 1, consistent with the overall decreasing profile, while the SU and DSU cluster report values of *r >* 1, consistent with the overall increasing pattern. The SU cluster furthermore reports stronger upregulation (*r* = 1.7) than the DSU cluster (*r* = 1.3). Peak expression timing also well reflect the pattern, with TU and SD cluster peaking early (4.3 hpf, 3.1 hpf) and the SU and DSU cluster reaching peak expression at much later developmental stages (83 hpf, 75 hpf).

Maternal degradation dynamics, characterised by the maternal half-life *t*_*m,half*_, likewise distinguish the clusters: the TU and SU cluster show fast maternal decay (0.71 h and 0.85 h), while SD and DSU cluster show slower maternal decay (1.3 h and 1.5 h).

The onset of zygotic genome activation, *t*_*zga*_ highlights the late activation of genes associated with the DSU cluster (*t*_*zga*_ = 16 *hpf*). Relative to this, activation times in the SD and SU cluster were estimated early at 2.5 hpf and 2.6 hpf, which aligns closely with the canonical timing of ZGA in zebrafish at approximately 3 hpf. The TU cluster instead shows an unusually early onset of transcription, with a cluster-center estimate of 0.97 hpf lying clearly before zebrafish ZGA. The timing of regulatory transition,*t*_*reg*_, shows a similar picture: early in the TU cluster (3.6 hpf), intermediate in the SD cluster (9.9 hpf), and latest in the SU (36 hpf) and DSU (47 hpf) clusters. Finally, the SD cluster is distinguished by high initial expression levels (33 TPM at cluster center) while the remaining clusters start at low to negligible expression levels (0.34 - 2.1 TPM).

We compared the cluster assignment changes between the shape-based (subscript S, in the following) and the parameter-based (subscript P) FCM clusters (Figure 9). The parameter-based *SD*_*P*_ cluster pre-dominantly consists of genes previously assigned to the *SD*_*S*_ cluster (81.94%), with smaller contributions from the *SU*_*S*_ cluster (12.86%) and the *DSU*_*S*_ cluster (5.20%). Similarly, *DSU*_*P*_ is largely composed of former *DSU*_*S*_ cluster genes (85.98%), with the remaining drawn from the *SU*_*S*_ (10.93%) and *SD*_*S*_ (3.09%) superclusters. In contrast, the *SU*_*P*_ cluster draws heavily from *DSU*_*S*_ genes (76.66%) compared to only 21.54% former *SU*_*S*_ and 1.70% *SD*_*S*_ genes. This also means that nearly half (49.94%) of the *DSU*_*S*_ cluster are now re-assigned to the cluster with earlier upregulation, and only 29.10% of *SU*_*S*_ cluster genes are reassigned to the corresponding parameter-based SU cluster. The newly identified *TU*_*P*_ supercluster is composed of 69.23% former *SD*_*S*_, 24.30% former *SU*_*S*_, and 6.42% former *DSU*_*S*_ genes, thus drawing primarily from genes following an overall downregulating pattern.

**Figure 9.**
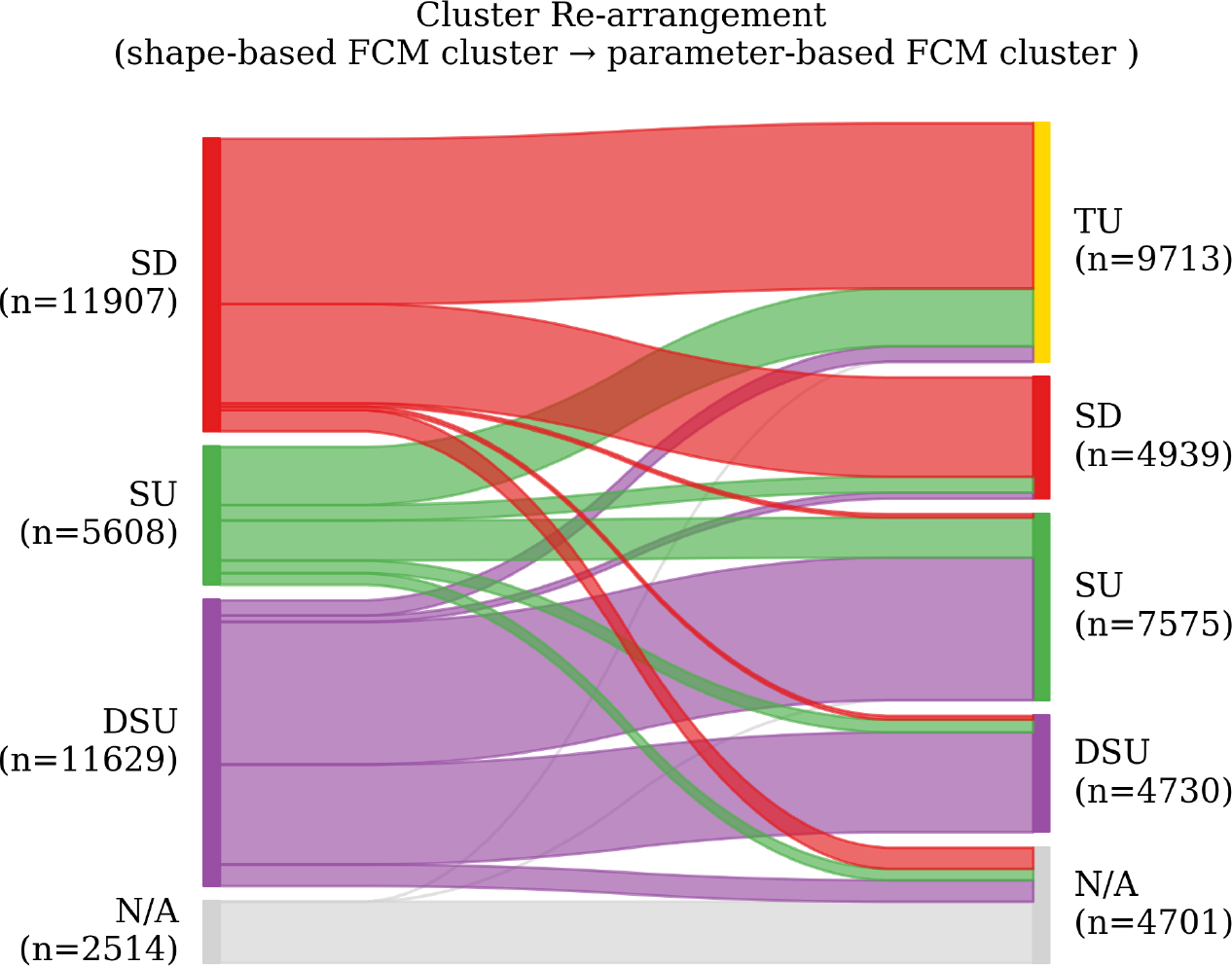
Genes changing cluster with using an GAM-based (left side) vs. mechanistic ODE modelling (right side) FCM clustering approach. The N/A cluster comprises the genes, that were excluded from the clustering analysis due to poor fit quality or expression levels below minimum threshold.

Alongside with the newly identified TU cluster, the composition of the SD cluster with respect to the decay-mode has shifted compared to the shape-based clustering. As described above, a substantial fraction of the *SD*_*S*_ cluster was reassigned to the *TU*_*P*_ cluster, leaving an intersection of 4,047 genes that remained in the SD cluster under both approaches (Fig. 10). Within the parameter-based SD cluster, the two decay modes are far more evenly distributed than before, with 48.1% of genes assigned as M-decay and 51.9% assigned as Z-decay, compared to the strong Z-decay dominance in the shape-based SD cluster.

Comparing the observed expression profiles (Fig. 10), the parameter-based SD cluster is also less dominated by pre-ZGA increases than its shape-based counterpart. While a pre-ZGA increase is still present, as visible in the mean expression trajectory, it is less pronounced (Δ_*t*=0,*t*=1.29_ = 0.26) than in the shape-based cluster Δ_*t*=0,*t*=1.29_ = 0.44). Together, these results indicate that the *MZ-Baseline* model is able to separate a subset of genes with a distinct, sharply transient expression pattern (TU) from genuinely maternally dominated expression (SD), a distinction the shape-based approach did not resolve.

In summary, the parameter-based approach identifies a distinct TU cluster characterised by a sharp, transient pre-ZGA increase in expression. The large reassignment of former SD-cluster genes to this cluster suggests that many genes originally grouped under sustained downregulation in fact follow this transient pattern, but that GAM-based smoothing underestimates this early dynamic.

**Figure 10.**
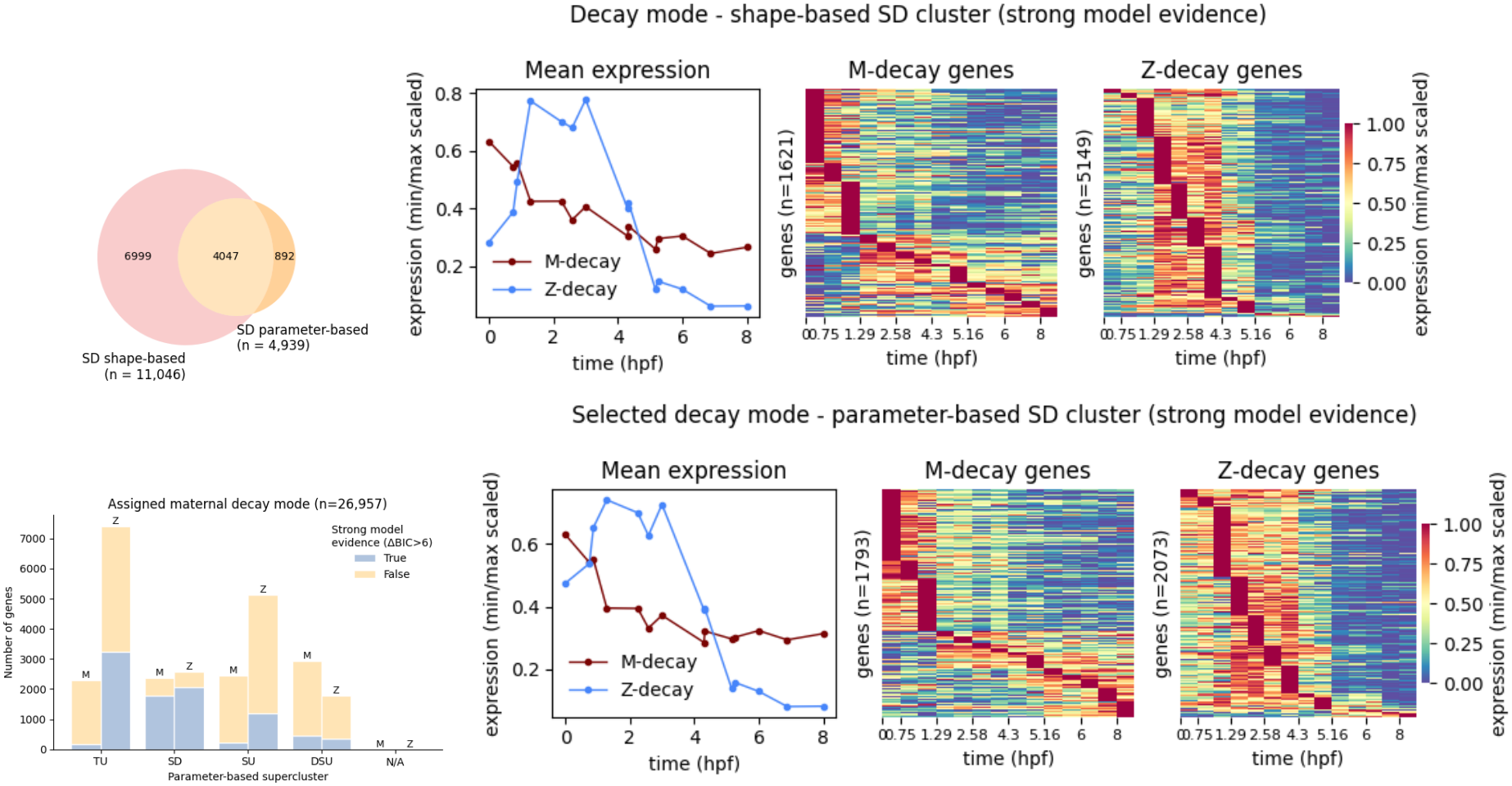
The Venn-diagram (upper left figure) shows the overlap of the shape-based and parameter-based SD cluster genes. The upper and lower middle figure shows the mean observed expression of genes from the shape-based and parameter-based SD-expression cluster, respectively, within an 8 hpf time-course, showing a strong model evidence for either the maternal (M) or zygotic (z) decay mode. M-decay genes show early degradation, starting within the first hour post-fertilisation, while for Z-decay genes, degradation only occur post ZGA, after approximately 3 hours post-fertilisation. The expression data was averaged across all sources and scaled by maximum expression. The heatmaps (right upper and lower figures) show the min/max-scaled expression profiles of the individual genes assigned to the preferred decay mode. M-decay genes show strong degradation before ZGA (3 hpf), while Z-decay genes are degraded post-ZGA. The lower left figure shows the breakdown of the preferred decay mode across parameter-based clusters, with blue bars indicating strong and beige bars indicating weak model evidence.

### 3.4 Evaluation of parameter-based FCM subclusters

The TU supercluster comprises two subclusters (Fig. 11), both characterised by sharp, early transient upregulation. TU-0 (n = 6,216) reaches peak expression before ZGA, while TU-1 (n = 3,497) increases more slowly. Intra-subcluster correlation (Fig. S10) confirms high pattern similarity within both subclusters (median Spearman’s *ρ* = 0.93 for TU-0, 0.89 for TU-1).

**Figure 11.**
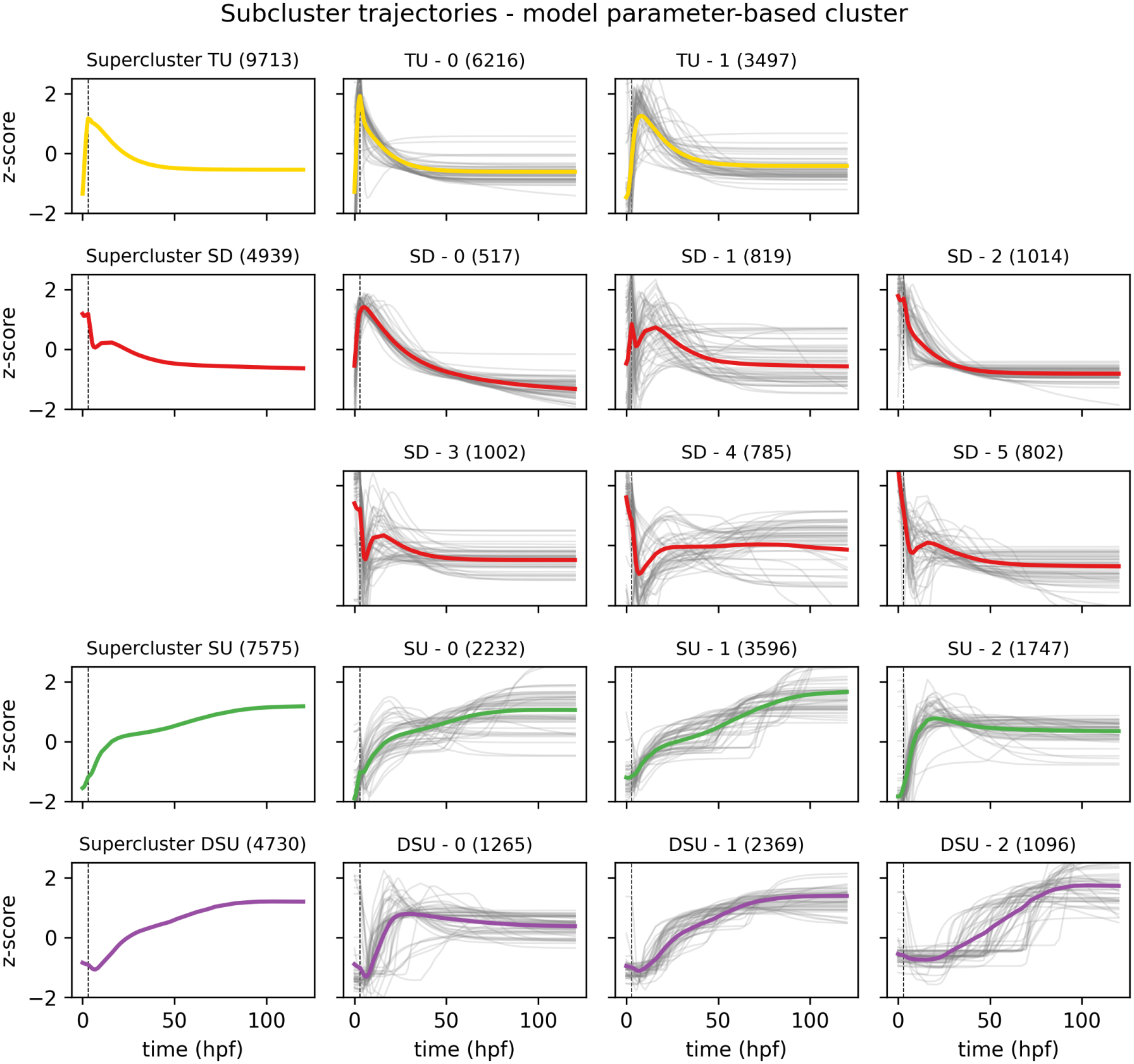
Parameter-based FCM subclusters. Colored lines show the average model trajectory per subcluster. Grey line show trajectories of single genes assigned to the subcluster.

The SD supercluster resolves into six subclusters with more heterogeneous patterns. SD-2 (1,014) shows consistently decreasing expression and the highest internal consistency of any SD subcluster (*ρ* = 0.99). Subclusters SD-3 (n = 1,002; *ρ*=0.73), SD-4 (n = 785) and SD-5 (n = 802; *ρ*=0.89), share a common early pattern with initially high expression levels, that decrease early, either before or during ZGA, and subsequently increase again. As indicated by the single-gene trajectories, SD-4 additionally contains genes that are up- or downregulated at later developmental times. This is also reflected by a poor, negative intra-cluster correlation of median *ρ* = −0,32. SD-1 is also a very heterogeneous subcluster (n = 819; *ρ* = 0.23), comprising a mix of profiles showing either transient upregulation or transient downregulation in early development. Notably, SD-0 (n = 517; *ρ*= 0.95) closely resembles a transient-upregulation pattern and, based on shape alone, might have been expected to fall within the TU supercluster rather than SD. The SU supercluster comprises three subclusters. SU-2 increases during ZGA and stabilises shortly after, with some trajectories showing a more transient peak. The overall correlation with the subcluster centroid however is good (n = 1,747; *ρ* = 0.78). Subclusters SU-0 (n = 2,232) and SU-2 (n = 3,596) both show steadily increasing expression with near-perfect correlation with their subcluster centroids (median *ρ* = 0.995). The two subclusters differ mainly in timing: SU-1 genes are characterised by delayed, post-ZGA increase, while SU-0 genes increase earlier.

The DSU supercluster likewise inherits three subclusters, that mainly differ by activation timing. DSU-0 (n = 1.265) shows a transient downregulation followed by a subsequent sustained post-ZGA increase from approximately 11 hpf. Overall intra-subcluster correlation is good with median *ρ*= 0.85. DSU-1 (n = 2,369) shows a similar but less pronounced initial downregulation, with expression levels increasing post-ZGA at around 14 hpf, later and more slowly than in DSU-0. Overall, DSU-1 has very high intra-subcluster correlation (*ρ*= 0.94) In DSU-2 (n = 1,096), activation is delayed considerably further, to approximately 37 hpf Intra-subcluster correlation is still good, but lower than in the other two DSU subclusters (*ρ*= 0.82), reflecting a broader distribution of activation and regulation times within this group (Fig. 12).

**Figure 12.**
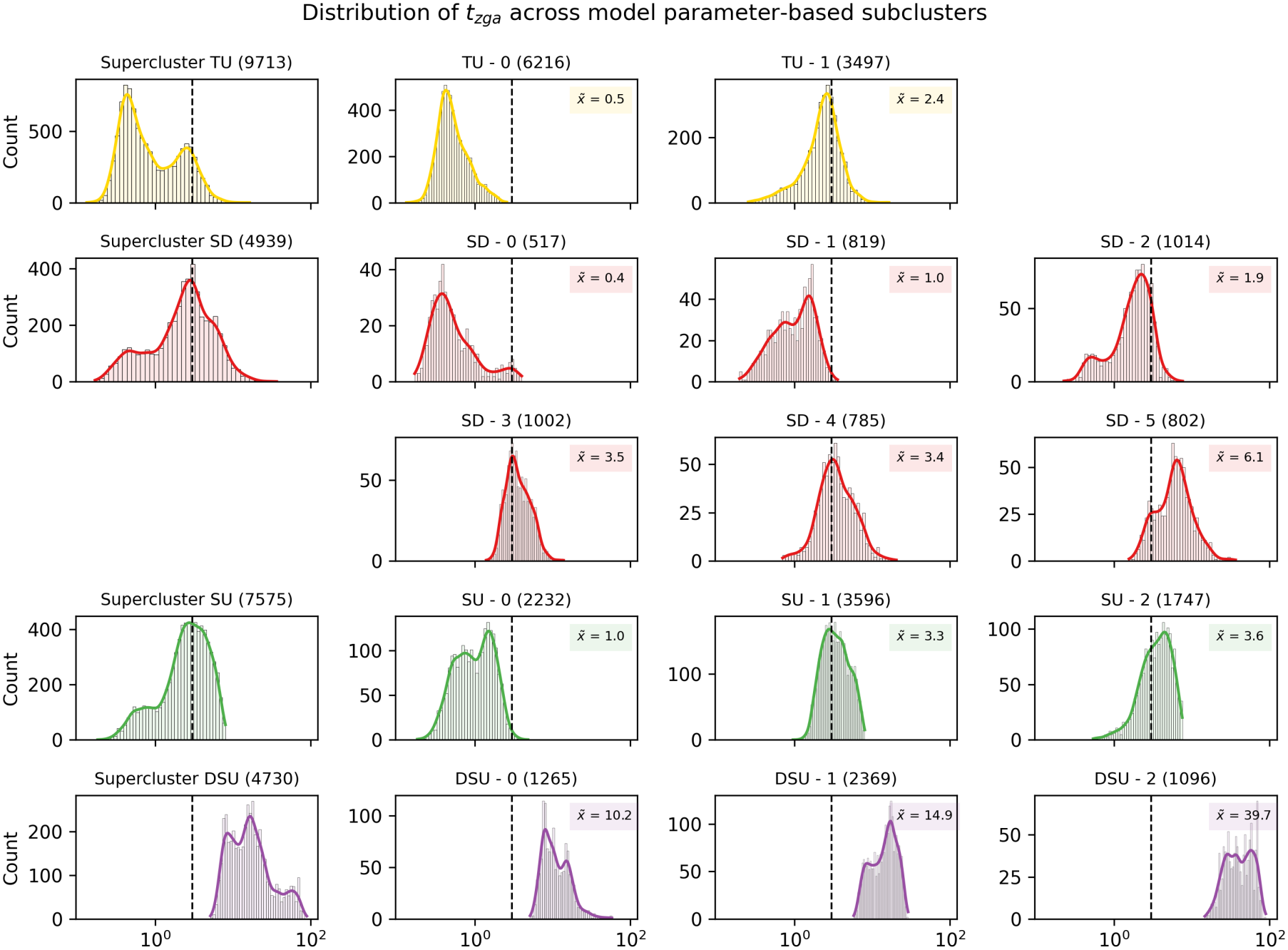
Distribution of ZGA onset (parameter *t*_*zga*_) across subclusters. The dashed line indicates the canonical timing of zebrafish ZGA at 3 hpf. Values before 3 hpf indicate early, pre-ZGA transcription. The label 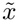 reports the median of the distribution per subcluster.

The activation time, estimated by model parameter *t*_*zga*_ deserves further attention. Expression profiles of TU cluster indicate very early zygotic transcription, lying before canonical timing of zebrafish ZGA of 3 hpf. Figure 12 show the distribution of *t*_*zga*_ estimates across subcluster. The zebrafish ZGA is displayed by the dashed line. The distribution confirms, that within the TU cluster, for largest fraction of genes (89.4%) the onset of ZGA was estimated before 3 hpf. Subcluster TU-0 consists entirely of pre-ZGA genes, with a median transcription onset of 0.5 hpf. TU-1 hold 70% genes activated before ZGA, with a median at 2.4 hpf. The SD cluster comprises subclusters with both primarily pre-ZGA onset (e.g. SD-0 with 97.1% and median *t*_*zga*_ = 0.4 hpf, SD-1 with 99.7% and median *t*_*zga*_ = 1.0 hpf) and primarily post-ZGA onset (SD-5 with 87% post-ZGA and median *t*_*zga*_ = 6.1 hpf). In the SU cluster 57.7% of genes show pre-ZGA increases, with SU-0 being the dominant pre-ZGA subcluster (99.4%, median *t*_*zga*_ = 1.0 hpf), while SU-1 and SU-2 show mixed contribution (41.9% and median *t*_*zga*_ = 3.5 hpf; 36.9%, and median *t*_*zga*_ = 6.1 hpf, respectively. In contrast, the DSU supercluster consist entirely of genes activated post-ZGA, with median *t*_*zga*_ estimates of 10.2 hpf, 14.9 hpf, and 39.7 hpf for DSU-0, DSU-1 and DSU-2, respectively.

## 4 Discussion

To investigate early embryonic gene expression dynamics, we compared two kinetic models: the *Basic* model and the *MZ-Baseline* model with variants M-decay and Z-decay for maternal dynamics and classified their dynamics with FCM-clustering (parameter-based-approach). The resulting pattern classification was compared to FCM-clustering on naive GAM-smoothed time-course expression trajectories (shape-based approach).

### 4.1 *MZ-Baseline* models outperform *Basic* transcription model

Each of the mechanistic model variants was fitted and evaluated using a zebrafish RNA-seq dataset comprising 31,658 genes across 120 hpf using a Bayesian inference approach, providing 55% of ZFE-genes with GOF-accepted mechanistic baseline models, which can be utilized in mechanistic ZFE models, e.g. in toxicokinetic models [28]. Despite higher complexity, the *MZ-Baseline* model substantially improved the accuracy of temporal expression pattern representation compared to the basic kinetic model (24% of genes fully GOF-accepted), and was preferred by BIC for the majority of genes.

This improvement can be explained by the structural extension of the *MZ-Baseline*. While the *Basic* model represents the most common formulation, assuming first-order decay [48, 49] and constant zygotic transcription [50, 51], the *MZ-Baseline* model extends this formulation by differentiating between maternal and zygotic transcripts and introducing a gene-specific activation time, consistent with the maternal-clock model of zygotic genome activation timing [52]. The model further incorporates a regulatory transition, which is expressed in either upregulation or downregulation of the transcript abundance. This allows modelling of transient expression peaks, a pattern frequently observed in early developmental time courses. This transcriptional regulation is modelled phenomenologically as a timing parameter, without considering the transcription factor network that causes these temporal dynamics. Despite this simplification, the *MZ-Baseline* model represents similar expression profiles as produced by feed-forward loops, without explicitly modelling the transcription factors.

The maternal state dynamic of the *MZ-Baseline* model differentiates between maternally (M-decay) and miR-430-mediated (Z-decay) degradation pathways. Analysis focused on early developmental stages confirmed that both decay mechanisms are pattern-wise prevalent among maternally provided genes. However, a key limitation of the two-state model evaluation is that the maternal and zygotic dynamics cannot independently be verified, as the provided dataset does not discriminate between maternal and zygotic transcripts. For genes with weak identifiability, the lacking differentiation between maternal and zygotic genes contributes directly to the convergence problems observed in a subset of model fits, since there is insufficient information in the data to confidently determine the maternal degradation rate *δ*_*m*_. Also, further mechanisms with a different timing of maternal clearance might be acting. Despite these limitations, the *MZ-Baseline* model proved to be a significant improvement compared to the *Basic* model for representing common baseline expression patterns in bulk RNA-sequencing data. The evaluation of the model fits showed that the *MZ-Baseline* model was generally preferred across all expression clusters, despite its higher complexity due to the larger number of free parameters.

### 4.2 FCM-clustering reveals strong discrepancy between shape-based and parameter-based approaches

The shape-based approach identified three major expression profiles: sustained downregulation (SD), sustained upregulation (SU) and delayed sustained upregulation (DSU). The parameter-based approach validated the finding of these same basic patterns, and additionally resolved a fourth supercluster, characterised by early transient upregulation (TU). Re-arrangement analysis showed that more than half of the genes were re-assigned from shape-based clusters to parameter-based clusters. Comparing these two clusterings showed, that the majority of former *SU*_*S*_ genes were re-assigned as *TU*_*P*_ genes, which could be represented particularly well by the *MZ-Baseline model*. This highlights the strength of the mechanistically informed parameter-based approach, as such pronounced early transient peaks were not detected using the smoothed trajectories.

It should be noted, however, that the GAM was fitted here in a deliberately naive way: no prior knowledge of the expected expression profile was provided, and the number of B-splines was fixed. The GAM could be used more effectively by incorporating the timing of zygotic genome activation (*t*_*zga*_ = 3 hpf) and placing a denser set of knots around it. This should allow the fit to follow the pre-ZGA increases seen in the data more closely. Additionally, the choice of fuzzy *c*-means as clustering method inherits some uncertainties that need to be addressed. FCM clustering defines clusters based on Euclidean distance measures and assigns each gene a “fuzzy” membership degree, which can be interpreted as a score of how strongly the gene relates to each cluster [31]. The fuzziness was not further considered in the clustering analysis, each gene was hard assigned to the cluster with the highest degree. FCM is dependent on a pre-defined number of clusters, which was chosen by Xie-Beni validity index [34]. Furthermore, the fuzziness parameter *m* is a sensitive parameter for FCM clustering, as it controls the overlap of clusters. The fixed values of *m* = 1.6 is within the commonly recommended range of [1.5, 2.5] [53], leaning towards harder cluster boarder. This value was not itself optimised, and a different choice of *m* could yield different cluster assignments or a different optimal cluster number under the XB index.

Apart from pattern identification, clustering offers an opportunity to improve prior definition of the temporal model parameters *t*_*zga*_ and *t*_*rep*_ as well as the steepness *s* of the activation curve. Under the assumption that genes from the same subclusters are activated or regulated at similar times, clustering could be used to define shared parameters. This could improve the parameter inference by using better-informed priors. Cluster based priors, based on purely maternal data (*δ*_*m*_) could be used to overcome the problem of convoluted maternal and zygotic transcription time-series, mentioned above. Especially noisy expression profiles, where no clear activation point is visible, would benefit from narrow priors. Furthermore, the model complexity could be reduced by fixing temporal parameters.

For less complex patterns as found in the DSU cluster, the *Basic* model proved to be suited for a large subset of genes. Thus, for describing baseline expression patterns of individual genes, the change in transcription rate based on temporal parameters is a good measure to capture the principal dynamics.

The *MZ-Baseline* models capture essential features of transcript production and degradation without requiring explicit representation of complex regulatory networks.

However, when investigating the effects of, for example, toxicant exposure, this simplified description may no longer be adequate. External stressors can induce a chain of reactions, affecting the expression levels of other genes. In such cases, the regulatory network of the target genes should be considered. There are common network motifs such as feed-forward loops, cascades or single-input modules that describe general dynamics in transcriptional regulatory networks [54]. Consequently, while the current modelling approach is appropriate for capturing baseline developmental dynamics, future extensions incorporating network information would be necessary to mechanistically interpret stress-induced transcriptional responses.

Finally, to increase biological interpretation of the clusters, a GO enrichment analysis to identify which biological functions, processes, or cellular components are significantly present, could be conducted.

### 4.3 Can poly-A tail binding explain sharp early-pre ZGA increases?

The *MZ-Baseline* model struggled most in the TU cluster (Fig. S9), as the sharp early pre-ZGA increase could not be captured accurately. This result suggests that the current models fail to account for certain dynamics present during the early developmental stages. As the zebrafish embryo is known to be transcriptionally silent until the zygotic genome activation at approximately 3 hpf, the observed pre-ZGA increases are unlikely to have a biological origin. A possible explanation for these pre-ZGA increases lies in the RNA-sequencing process. The RNA-seq dataset was generated using polyA+ selection as library preparation method [29, 30]. The library preparation for RNA sequencing using oligo(dT) based selection methods to isolate mRNAs for the total RNA sample introduces a bias towards transcripts with longer poly(A) tails, as short tails bind oligo(dT) beads less efficiently. Consequently, the abundance of short-tailed mRNAs in RNA-seq samples where oligo(dT) capture was used is underestimated [55, 56]. The poly(A) tail length is strongly coupled to mRNA stability and translation efficiency, thus the length is dynamically regulated during embryogenesis. Maternal mRNAs are usually deposited in the embryo with short tails, which leaves them in a dormant state and enables their storage. After the first few cleavages, maternal mRNA undergoes cytoplasmic polyadenylation (CPA), which elongates the poly(A) tail to increase the translation efficiency [15]. This growth in tail length also increases the oligo(dT) binding affinity during the RNA-seq library preparation, resulting in an apparent pre-ZGA increase in transcript abundances. Such technical biases are currently not considered by the *MZ-Baseline* model. Adding a poly(A)-aware extension could improve fit quality and biological interpretability.

Furthermore, apparent increases in transcript abundance could also result from differential transcript stability during early developmental stages. This effect would be a consequence of the TPM normalisation. When unstable transcripts degrade rapidly during the early stages, while the embryo is still transcriptionally silent, the TPM of stable transcripts increases in relation to unstable ones. Re-examining the raw count data could help to disentangle these normalisation artefacts from polyA+ biased increases.

## Acknowledgements

Computations on the HPC of Osnabrück University were funded by the Deutsche Forschungsgemeinschaft (DFG, German Research Foundation) - 456666331.

## Supporting information

### S1 Data and code availability

The expression dataset and FCM clustering can be accessed via Github:

https://github.com/mariegrho/FCM_clustering.git the code used for the Baseline model can be found under https://github.com/mariegrho/Expression_baseline_models/tree/hpc_fits

### S2 Goodness-of-fit metrics

Different types goodness of fit metrics were used to evaluate the pre-gene model fit quality from the ArviZ InferenceData.

#### NUTS-specific convergence diagnostics per parameter (posterior, sample stats)

The R-hat 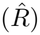 convergence diagnostic evaluates whether the four MCMC chains have converged to the same target distribution. 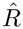 value close to 1.0 indicates the the Markov-Chains have mixed well. Here, we used the threshold 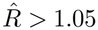 as indicator for good convergence. The metric was drawn from *arviz stats.rhat*.

Furthermore, the bulk effective sample size (bulk-ESS) measures the sampling efficiency for the center of a posterior distribution. Generally, a higher ESS relates to better convergences. A minimum threshold of ESS *<* 400 (100 per chain) was used as additional convergence criteria. Here, the *ArviZ* integrated function *arviz stats.ess* was used.

#### Bayesian metrics (Log-Likelihood), comparison of model alternatives

The Bayesian Information Criterion (BIC) is used to compare multiple models fitted to the same dataset. It balances model fit and complexity by incorporating both the maximum log-likelihood 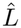, the number of free parameters *k* and the number of observations *n*. A lower BIC value indicates a better model as it suggests a superior balance between goodness of fit and parsimony.

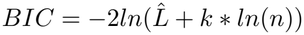

In terms of BIC comparison between two models, a Δ*BIC >* 6 is interpreted as strong evidence towards the model with lower BIC [57].

#### Posterior predictive coverage (PPV) (posterior predictive)

Posterior predictive coverage measures the fraction of observed points falling within their 50% and 90% posterior predictive interval. A systematic over- or under-coverage flags model misspecification or a noise model that’s too narrow/wide.

#### Point-estimate fit quality (posterior model fit, observed data)

The Normalised Root Mean Squared Error (NRMSE) expresses the model’s prediction error normalised by the range of the observed data, allowing comparison across datasets with different scales [58]. The NRMSE is applied to the untransformed dataset and calculated by:

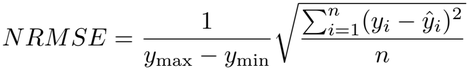

where *y*_*i*_ and *ŷ*_*i*_ represent the observed and predicted values, respectively and *n* is the total number of observations. Lower NRMSE values indicate better model performance. Here, an NRMSE *<* 0.3 is defined as a good-fit threshold.

Spearman’s rank-order correlation coefficient *ρ* quantifies the strength and direction of the monotonic relationship between the two datasets and serves thus as a similarity measure [59]. It is defined as:

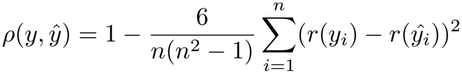

where n represents the number of observation points and *r*(*y*_*i*_) is the rank of *y*_*i*_ in the expression profile of gene *y*. Values close to +1 indicate a strong positive monotonic relationship, meaning that as *y* increases, so does *ŷ*, whereas values near −1 indicate an opposing behaviour. The threshold for good pattern accuracy was defined as *ρ >* 0.5. It was implemented using the *scipy.stats*-method *spearmanr* [60].

### S3 Model parameter

Table S1 summarises the model parameters and initial values.

**Table S1.**
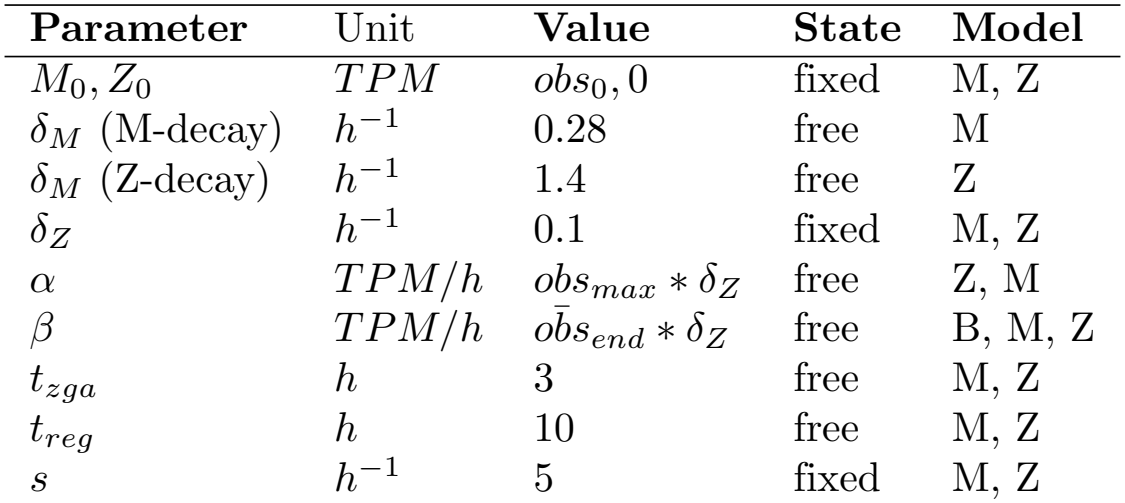
Prior scales for model variants. Priors are defined with a lognormal distribution. Model abbreviations: B = Basic, M = MZ-Baseline M-decay and Z = MZ-Baseline Z-decay

| **Parameter** | Unit | **Value** | **State** | **Model** |
|---|---|---|---|---|
| M<sub>0</sub>, Z<sub>0</sub> | TPM | obs<sub>0</sub>, 0 | fixed | M, Z |
| δ<sub>M</sub> (M-decay) | h<sup>-1</sup> | 0.28 | free | M |
| δ<sub>M</sub> (Z-decay) | h<sup>-1</sup> | 1.4 | free | Z |
| δ<sub>Z</sub> | h<sup>-1</sup> | 0.1 | fixed | M, Z |
| α | TPM/h | obs<sub>max</sub> * δ<sub>Z</sub> | free | Z, M |
| β | TPM/h | $o\bar{b}s_{end} * \delta_Z$ | free | B, M, Z |
| t<sub>zga</sub> | h | 3 | free | M, Z |
| t<sub>reg</sub> | h | 10 | free | M, Z |
| s | h<sup>-1</sup> | 5 | fixed | M, Z |

### S4 Shape-based FCM clustering

For the clustering analysis, only GAMs-based trajectories that passed the quantitative (NRMSE¡0.25) or qualitative (Spearman’r *ρ*¿0.5) fit-quality metrics were selected for the clustering analysis.

The GAMs approach provided an overall good quantitative fit, with an median NRMSE of 0.13 and an acceptance rate of 99.2% across all genes. A positive pattern correlation was achieved for 87% of genes, with an median *ρ* of 0.82.

Based on the maximum expression value across sources, 2,350 genes were rejected from the clustering analysis, showing only marginal expression below 0.1 TPM over the complete time-course.

In total, 29,144 GAMs fits passed the goodness-of-fit criteria and were further analyised in the clustering analysis.

**Figure S1.**
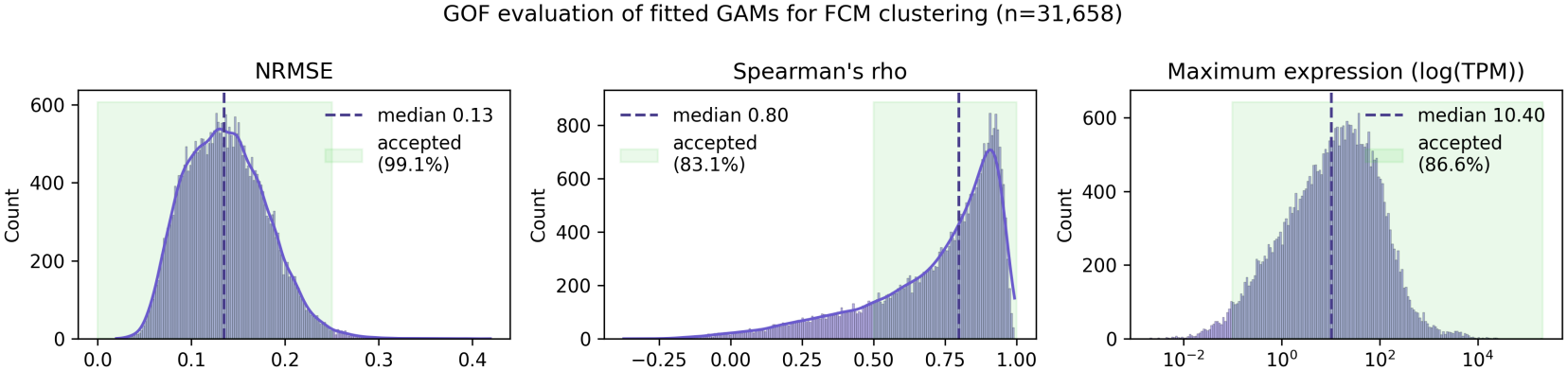
Distribution of NRMSE and *ρ* values of GAMs fits.

Figure S2 shows the selection metrics for the number *c* of fuzzy c-means clusters over a range of c=2 to c=9 using Xie-and-Beni index and the stability score as validity measure. The optimal number of clusters was determined by the minimun XB-index.

Figure S3 show the cluster centers of shape-based FCM superclusters and subclusters, alongside with 50 fitted smooth trajectories that were assigned each subcluster.

### S5 Model fit evaluation

Figure S5 shows the proportion of accepted ODE model fits across shape-based subclusters. Proportions were calculated based on the total non-zero expression dataset (n=31,658 genes). Genes were flagged as “accepted”, when the model fit passed either the quantitative or qualitative GOF criteria.

Figure S6 shows the goodness-of-fit evaluation of the *MZ-Baseline* and *Basic* model fits.

### S6 Parameter-based FCM clustering

For parameter-based FCM clustering, the *MZ-Baseline* model fits were evaluated based on quantitative and qualitative fit accuracy. In total, 26,957 genes passed the low-expression filter and GOF criteria for the *MZ-Baseline* model and were subsequently used for the clustering analysis.

Prior to applying FCM clustering, the fitted model parameters were log-transformed and z-scored to eliminate the orders of magnitude across genes The Figure S7 shows the distribution of the scaled parameter values, that were used as database for the parameter-based fuzzy c-means clustering analysis.

Figure S8 shows the selection metrics for the number *c* of fuzzy c-means clusters over a range of c=2 to c=9 using Xie-and-Beni index and the stability score as validity measure. The optimal number of clusters was determined by the minimun XB-index.

Figure S11 shows the mean observed expression values over time, partitioned in shape-based super-clusters (left figure) and parameter-based supercluster (right figure).

**Figure S2.**
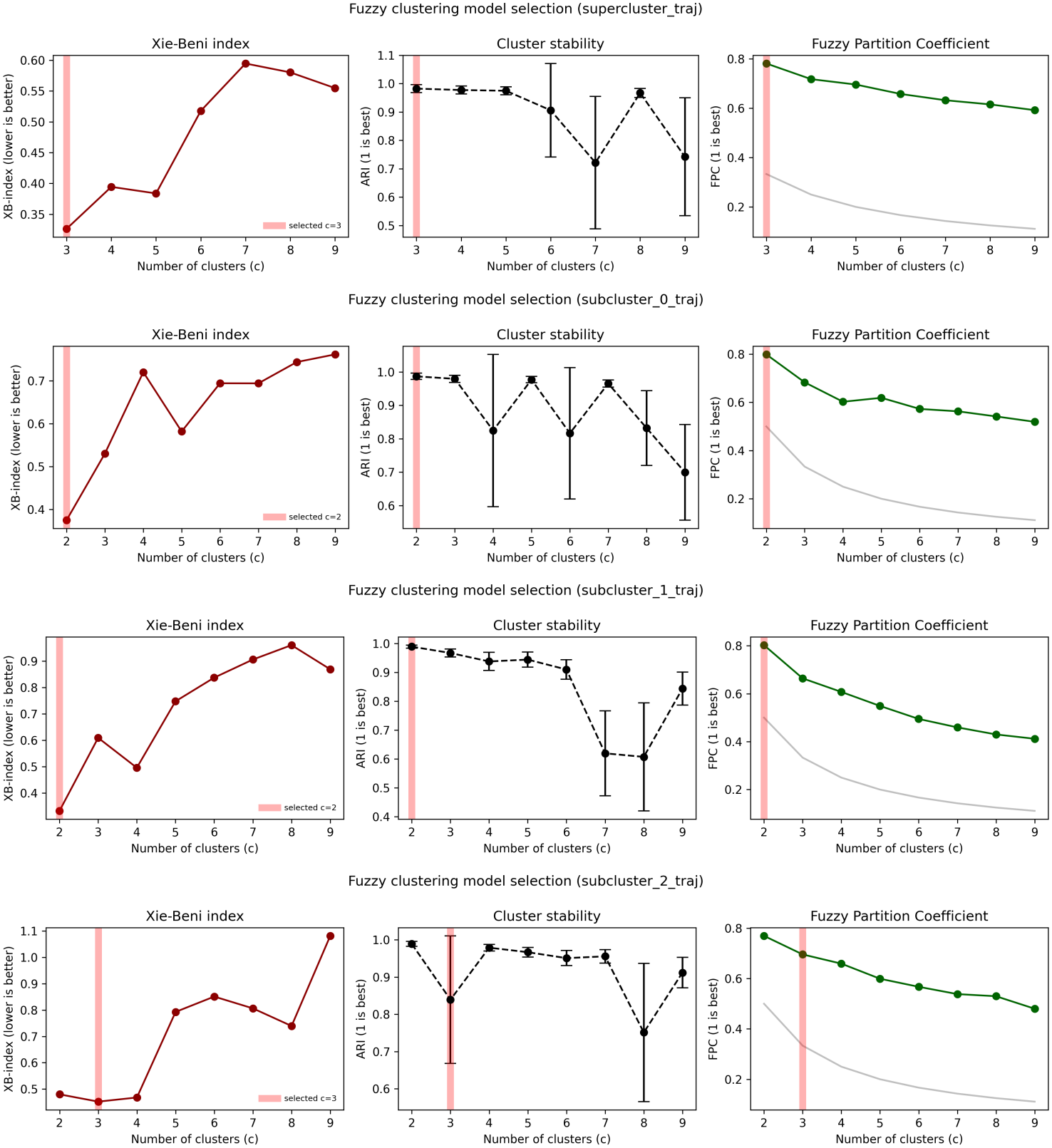
Xie-and-Beni index for FCM cluster selection alongside with the stability score and mean FPC for shape-based superclusters and corresponding subclusters. The red lines shows the selected number of clusters c based in the XB-index.

**Figure S3.**
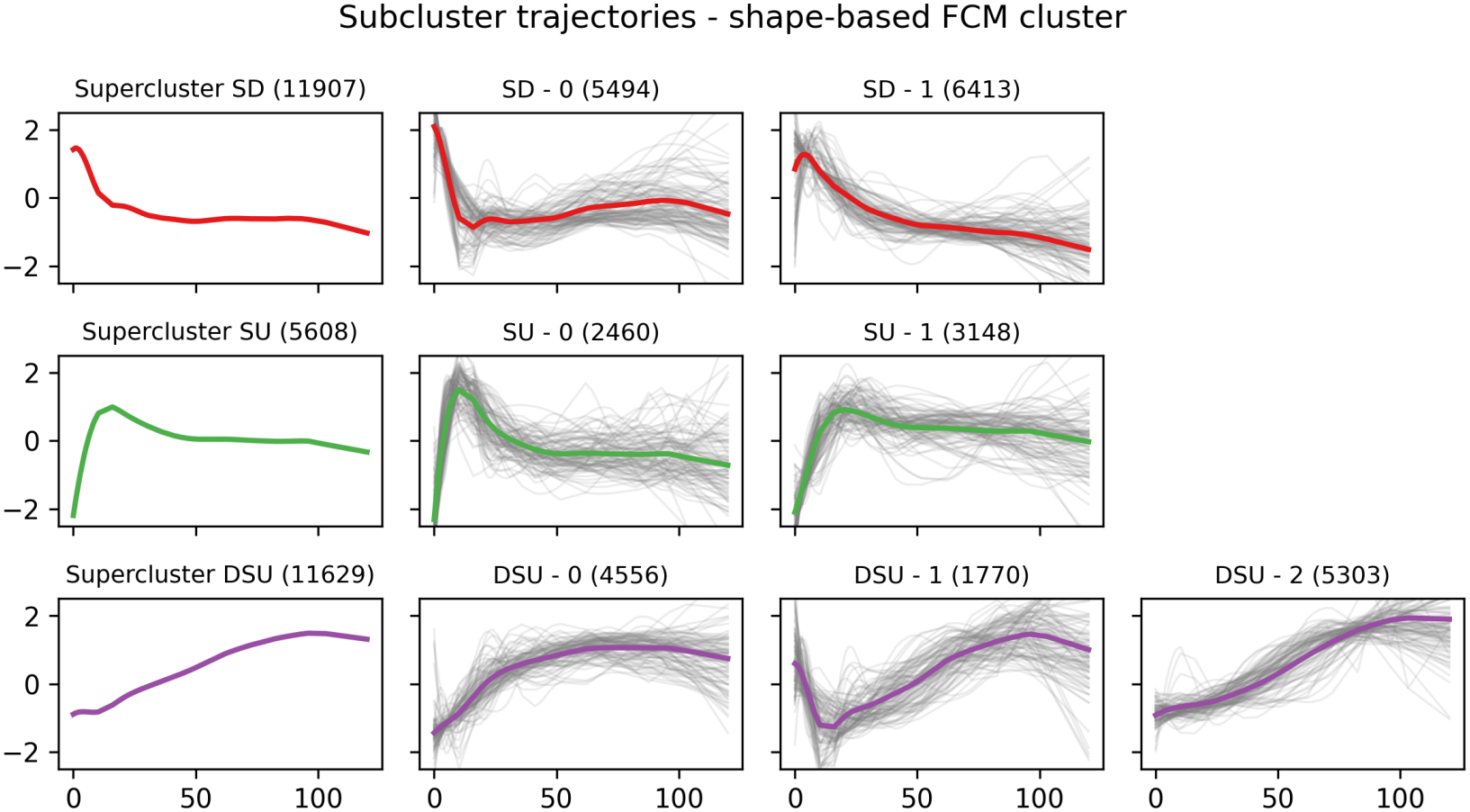
Average expression per shape-based subcluster (bold colored lines) alongside with the fitted GAMs trajectories of a random subset of the subcluster (thin grey lines). Expression values are log2(TPM+1) transformed and standardised by z-score.

**Figure S4.**
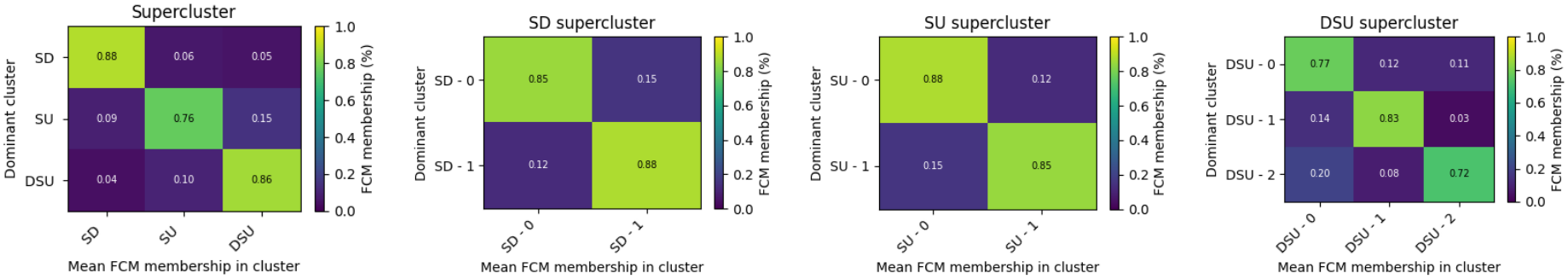
Mean FCM membership per shape-based supercluster and subcluster. An FCM membership of 1 refers to no-overlap with other clusters.

**Figure S5.**
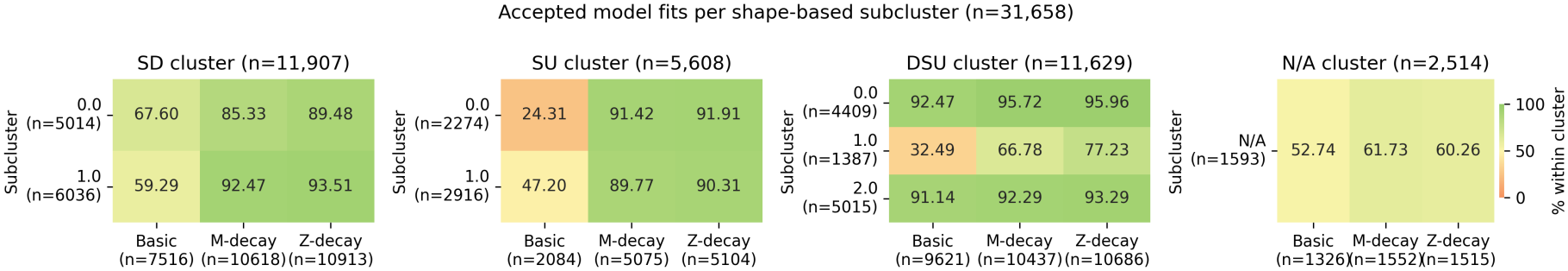
Aggregated results of “fully accepted” baseline model fit per shape-based subcluster.

**Figure S6.**
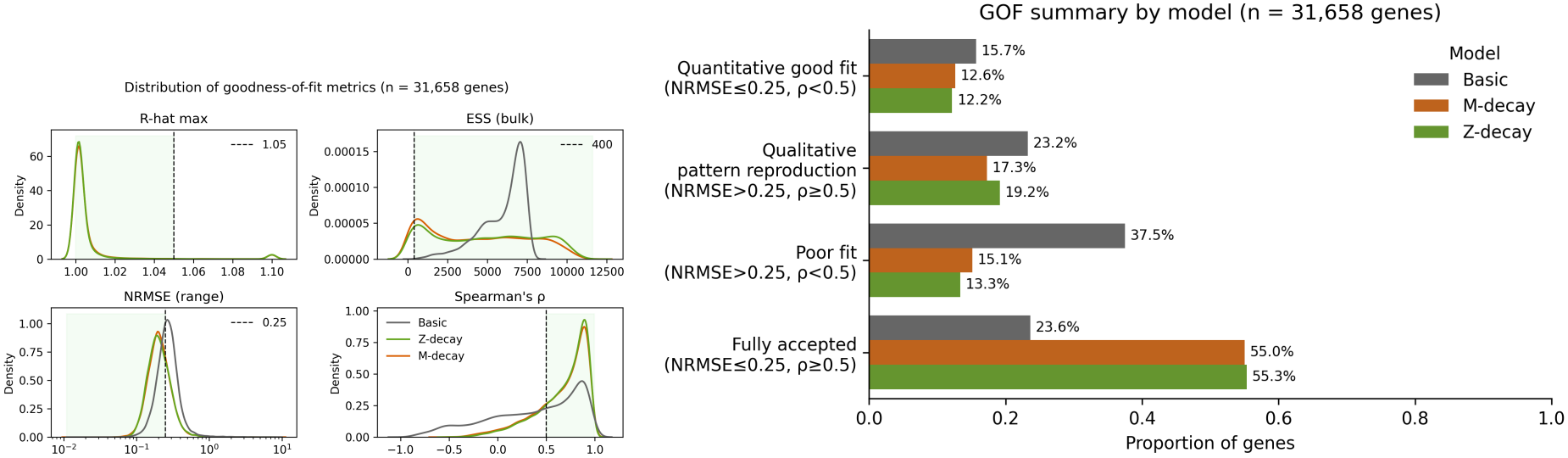
Goodness of fit summary of the baseline model fits per model. Left figure: Distribution of the goodness-of-fit metrics across the three model variants. Right figure: Summary of the model fit metrics. The following criteria were applied: “Poor fit”: all good fit criteria failed (*NRMSE >* 0.3, *ρ <* 0.5,), “Qualitative pattern”: (*NRMSE >* 0.3, *ρ >* 0.5,), “Quantitative fit”: (*NRMSE <* 0.3, *ρ <* 0.5,), “Fully accepted”: all good fit criteria are fulfilled (*NRMSE <* 0.3, *ρ >* 0.5,).

**Figure S7.**
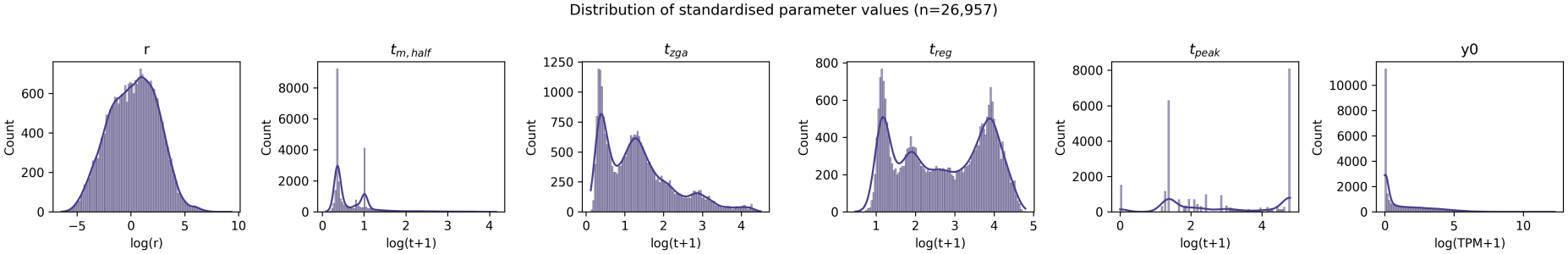
Distribution of log-scaled and z-scored parameter values, that were used for ODE parameter based FCM clustering.

**Table S2.** Intra-subcluster correlation of parameter-based FMC clusters, assessed by Spearman’s *ρ*.

|  |  | mean | median | std | count |
|---|---|---|---|---|---|
| supercluster | subcluster |  |  |  |  |
| TU | 0 | 0.781 | 0.928 | 0.444 | 6216 |
|  | 1 | 0.765 | 0.887 | 0.331 | 3497 |
| SD | 0 | 0.932 | 0.950 | 0.088 | 517 |
|  | 1 | 0.221 | 0.215 | 0.503 | 819 |
|  | 2 | 0.903 | 0.993 | 0.241 | 1014 |
|  | 3 | 0.402 | 0.584 | 0.457 | 1002 |
|  | 4 | -0.135 | -0.323 | 0.496 | 785 |
|  | 5 | 0.659 | 0.867 | 0.406 | 802 |
| SU | 0 | 0.889 | 0.995 | 0.224 | 2232 |
|  | 1 | 0.966 | 0.995 | 0.097 | 3596 |
|  | 2 | 0.709 | 0.782 | 0.297 | 1747 |
| DSU | 0 | 0.737 | 0.845 | 0.323 | 1265 |
|  | 1 | 0.926 | 0.935 | 0.083 | 2369 |
|  | 2 | 0.730 | 0.821 | 0.198 | 1096 |

**Figure S8.**
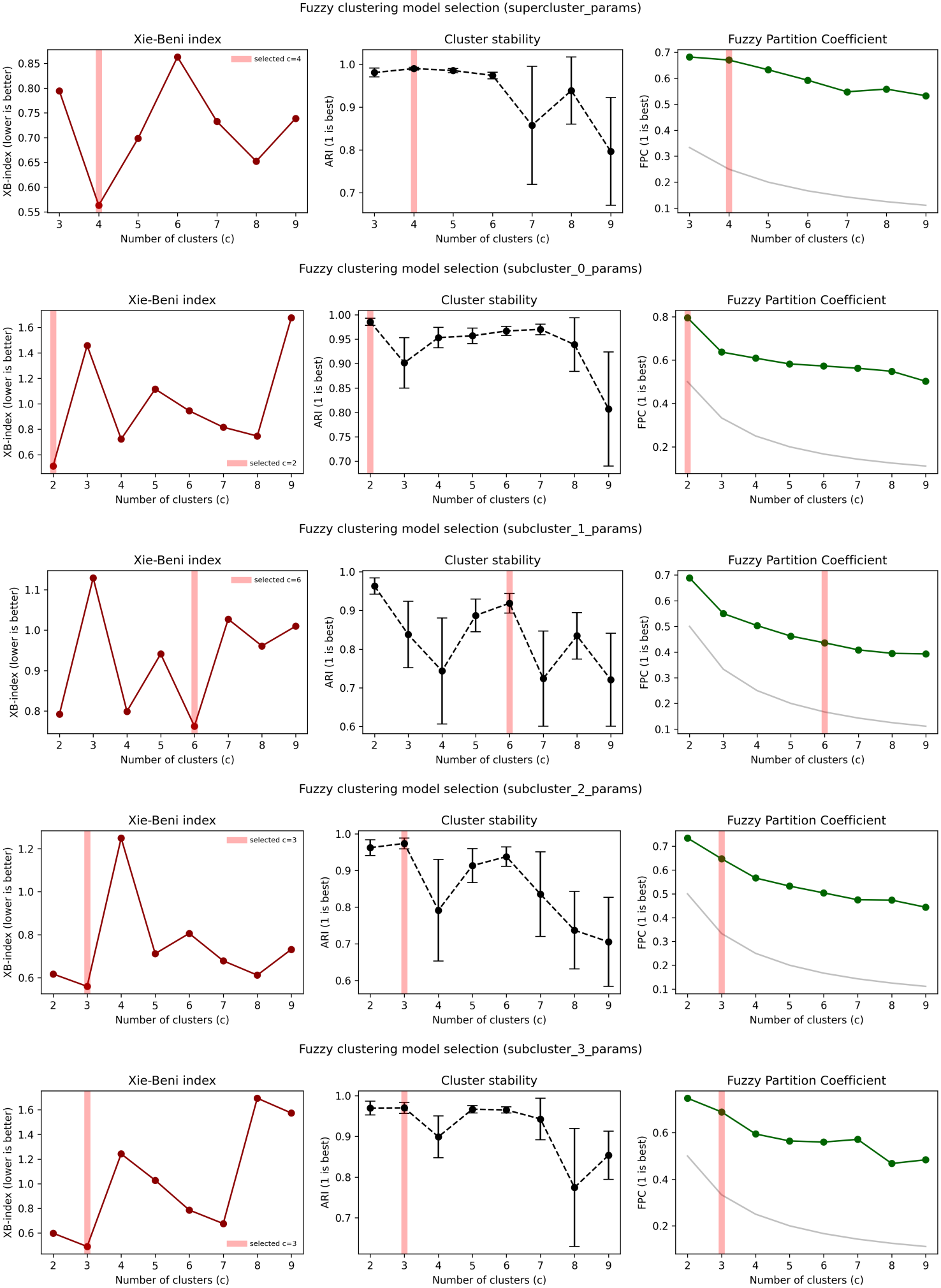
Xie-and-Beni index for FCM cluster selection alongside with the stability score and mean FPC for parameter-based superclusters and corresponding subclusters. The red lines shows the selected number of clusters c based in the XB-index.

**Figure S9.**
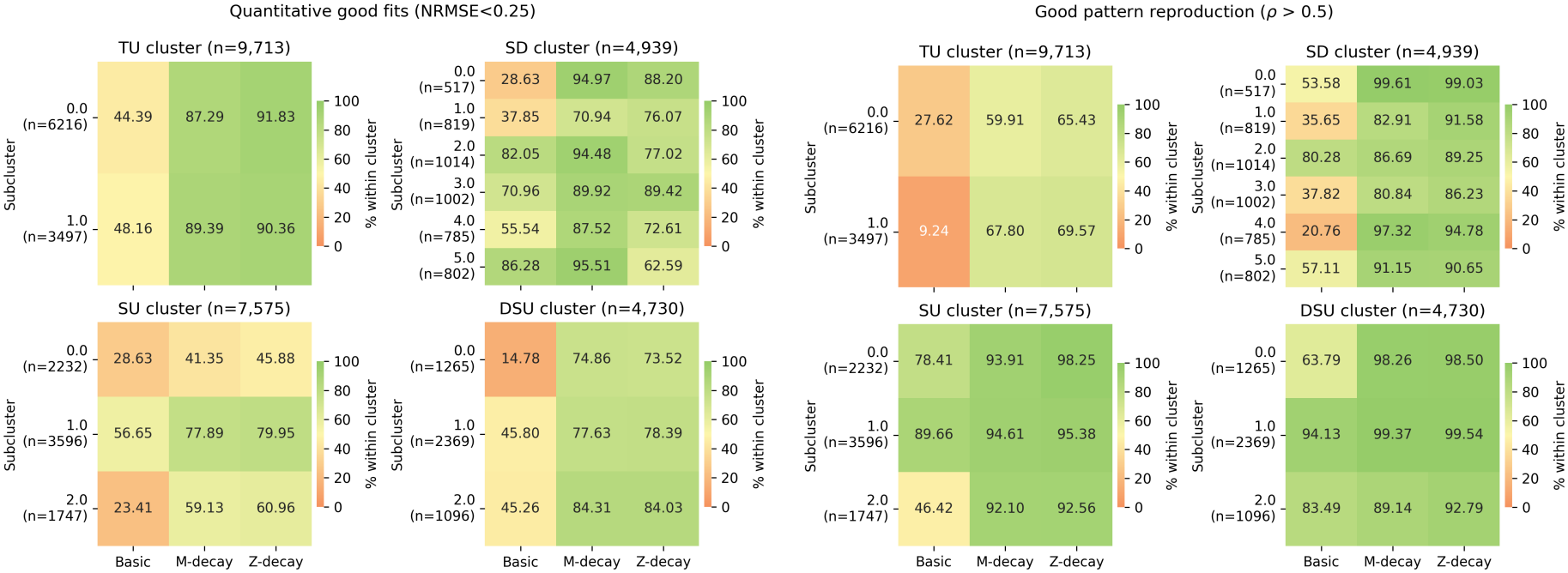
Quantitative and qualitative fit quality of model-fit across parameter-based subclusters.

**Figure S10.**
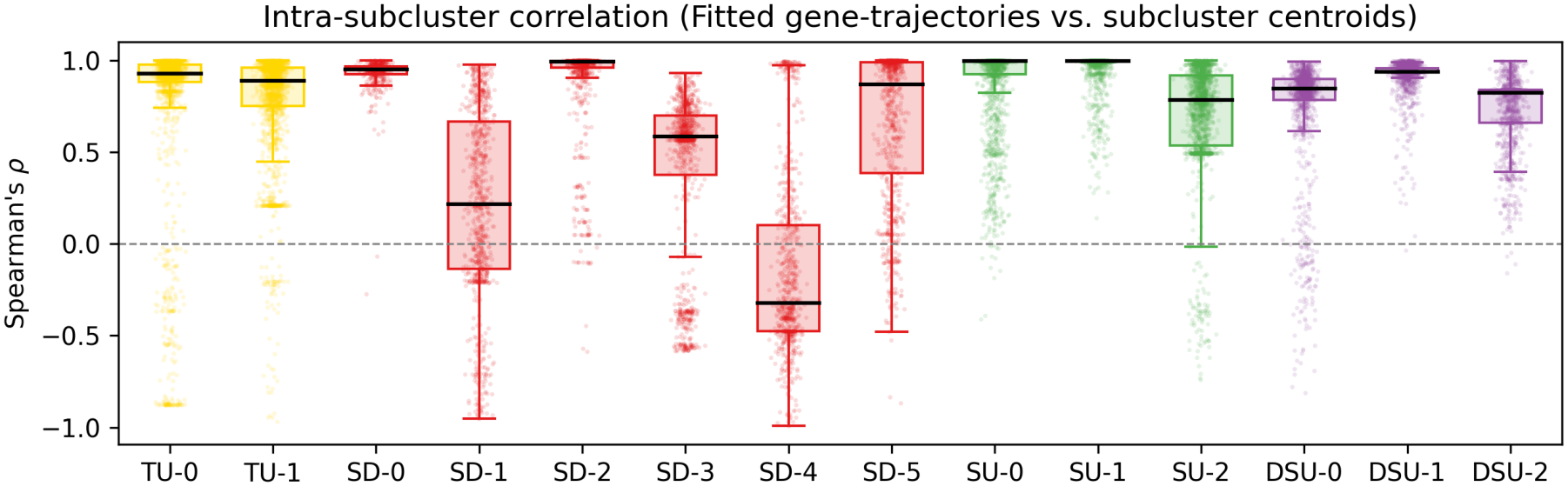
The boxplot shows the intra-subcluster correlation of fitted model trajectories to their subcluster centroid. Correlation was assessed by Spearman’s *ρ*. See Table S2 for values.

**Figure S11.**
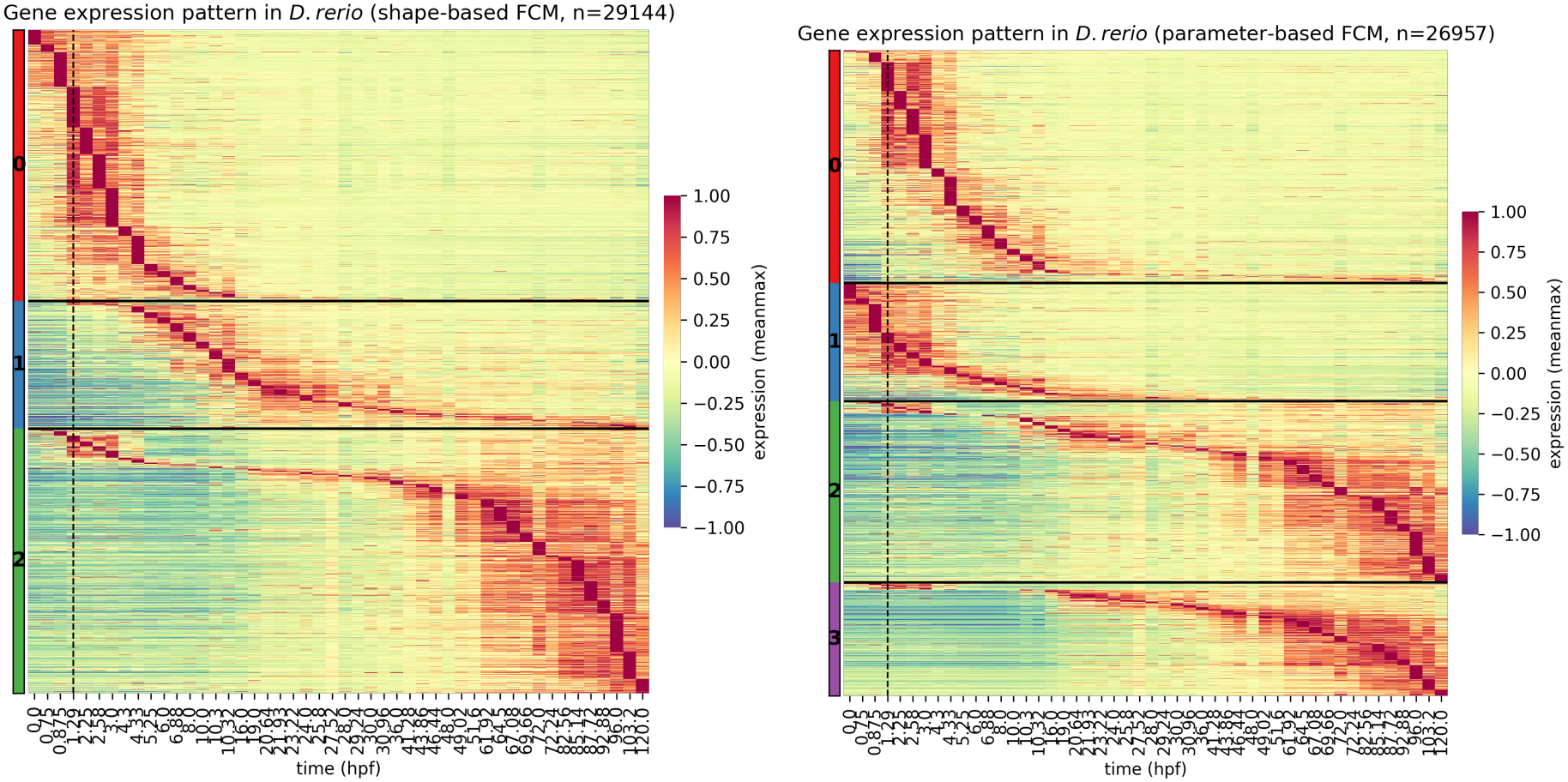
Heatmap of mean temporal gene expression, ordered by shape-based FCM clusters (left) and paramerer-based FCM cluster (right). Genes are sorted by increasing maximum expression, TPM expression values are min/max normalised.

